# Metabolite patterns in human myeloid hematopoiesis result from lineage-dependent active metabolic pathways

**DOI:** 10.1101/2020.07.09.195156

**Authors:** Lars Kaiser, Helga Weinschrott, Isabel Quint, Folker Wenzel, Markus Blaess, Manfred Jung, Matthias Kohl, Hans-Peter Deigner

## Abstract

Assessment of hematotoxicity from environmental or xenobiotic compounds is of notable interest and is frequently assessed via the colony forming unit (CFU) assay. Identification of the mode of action of single compounds is of further interest, as such often enables transfer of results across different tissues and compounds. Metabolomics displays one promising approach for identifying such, nevertheless, suitability with current protocols is restricted. Here, we combined an HSPC expansion approach with distinct lineage differentiations, resulting in formation of erythrocytes, dendritic cells and neutrophils. We examined the unique combination of fluxes in glycolysis, glutaminolysis, polyamine synthesis, fatty acid oxidation and synthesis, as well as glycerophospholipid and sphingolipid metabolism. We further assessed their interconnections and essentialness for each lineage formation. By this, we provide further insights into metabolic fluxes during differentiation of HSPC into different lineages, enabling profound understanding of possible metabolic changes in each lineage caused by exogenous compounds.

## Introduction

Assessment of toxic properties from environmental or xenobiotic compounds remains a major field of research. Mammalian laboratory animals are currently considered as the ‘gold standard’ in toxicology.^1^ However, as mammalian models are expensive and time-consuming, several *in-vitro* models have been developed and applied.^2–4^ Indeed, models based on primary human cells are currently believed to more accurately reflect *in-vivo* responses towards selected compounds.^1^ Furthermore, current research sets a special focus on primitive cell types (e.g. embryonal stem cells, induced pluripotent cells), as these are appreciated to be particularly vulnerable towards exogenous stimuli and also display a model for developmental toxicity testing.^5–7^ In this regard, models based on hematopoietic stem and progenitor cells (HSPC) are often used, to assess possible hematotoxicity of compounds. Assessment of hematotoxicity in fact is of notable interest, as hematopoiesis continuously occurs during whole individuals lifetime and large quantities of all blood cells arises daily from hematopoietic stem cells. Furthermore, as several environmental contaminants are known to cross the placenta, fetal HSPC are of particular interest, as these display potential sentinels of later-life hematopoietic disorders.^8^ To date, traditional hematotoxicity testing is performed using colony forming unit (CFU) assays, however, also more lineage-specific models have recently been developed.^9,10^

Identification of the toxicological mode of action of single compounds is of particular interest, as such often enables transferability of results across different tissues sharing similar metabolic characteristics. At current, omics technologies are promising for identifying such features in common, as these enable rapid screening.^11^ Indeed, metabolomics presumably displays the most promising approach, as it is thought to display the phenotype most accurately and enables prediction of toxic effects in very early stages.^12^ A significant flaw of metabolomics, however, is the need of large amounts of sample material (typically >1×10^6^ cells per replicate), making metabolomics mostly incompatible for large screening approaches with the afore mentioned hematopoietic models.^13–16^ Furthermore, profound knowledge of the total cellular metabolism in the different lineages is a prerequisite for interpretation of observed metabolic changes, as it ultimately defines the response of cells towards different stimuli.^17,18^ As example, intracellular levels of NADPH define the response towards reactive oxygen species (ROS)-inducing compounds. Intracellular levels of NADPH in turn, are dependent of pentose phosphate pathway (PPP) activity, as well as of activity of *IDH1* and *ME1*.^19,20^ Thus, two cell types differing in their PPP activity may exhibit distinct responses towards the same compound, rendering profound knowledge of active metabolic pathways an essential feature for understanding. In context to hematotoxicity, cell type specific effects are already well appreciated, several compounds are known to induce lineage-specific effects. For example, erythroid progenitors are known to be more susceptible against lead, benzene or N-acetylcysteine than other lineages.^9,21–23^ In case of 3’-azido-3’-deoxthymidine (Azidothymidin), this cell type specific effect is even more pronounced; erythroid progenitors are substantially reduced, while granulocyte/macrophage as well as megakaryocytic progenitors remain unaffected.^24^ Nevertheless, it also should be noted, that also endogenous compounds may possess an impact on hematopoiesis. For example, lactate was recently shown to promote erythropoiesis via induction of ROS.^25^ Such effects may be identified by usage of classical colony forming unit (CFU) assays or more recently developed hematopoietic differentiation models.^10^ For elucidation of the mode of action behind such effects, however, profound knowledge of the similarities and differences in the metabolism of each lineage is essential. Likewise, identification of relations between active metabolic pathways and specific responses likely enables prediction of similar response patterns on other compounds with analogical modes of actions. Moreover, such relations may also enable prediction of response patterns across different tissues, leading to a better prediction of possible, tissue specific, toxic effects during drug development or testing of xenobiotics.

Indeed, lineage-dependent regulatory involvement of single metabolic pathway activity during hematopoiesis is quite evident. Regulation of fatty acid oxidation (FAO) for instance, seems to be crucial for hematopoietic stem cell (HSC) maintenance, since blocking of FAO promotes HSC commitment.^26^ However, autophagy-mediated generation of free fatty acids and subsequent degradation via FAO is crucial for neutrophil differentiation, indicating active FAO during differentiation of (at least) some lineages.^27^ Furthermore, lymphocytes, neutrophils and macrophages utilize glutamine at high rates under catabolic conditions (e.g. sepsis), underlining the importance of glutaminolysis during HSPC differentiation.^28^ Blocking glutaminolysis in erythropoietin (EPO)-stimulated HSPC, however, leads to a shift from erythroid commitment towards a myelomonocytic fate.^29^ Therefore, modulation of glutaminolysis by xenobiotic compounds may also result in lineage-specific toxicity. Nevertheless, the assumption that glutaminolysis solely defines erythroid lineage commitment falls quite short, since it has been shown recently that blocking choline generation from phosphatidylcholine also impairs erythroid differentiation.^30^ The role of phosphatidylcholine degradation within differentiation of other myeloid lineages, however, remains vague. In addition, several studies suggest a relation of polyamines with erythroid differentiation, their role in other lineages, however, again remains inconclusive.^31–33^ Taken together, the essentialness of several different metabolic pathways during defined HSPC differentiation has already been shown for selected lineages. The general activity, interconnections and fluxes between the different metabolic pathways, also within other lineages, however, remains still unclear. Therefore, a direct comparison of metabolic fluxes within different hematopoietic lineages is desirable, in order to further elucidate the mode of action behind possible lineage specific effects.

Here we combined a published HSPC expansion approach with distinct lineage differentiations from the literature, resulting in formation of erythrocytes, dendritic cells (DC) and neutrophils. Due to the initial expansion step, large cell numbers can be generated with this approach, making it highly suitable for omics-based toxicity testing. Further assessment of metabolic and transcriptional changes during lineage formation resulted in unique and common metabolite sets, reflecting distinct metabolic changes in several interconnected pathways (namely glycolysis, glutaminolysis, polyamine synthesis, fatty acid oxidation and synthesis, as well as glycerophospholipid and sphingolipid metabolism). We further assessed the essentialness of glutaminolysis, polyamine synthesis and FAO for differentiation of each lineage, confirming the proposed fluxes. While several pathways were active in different lineages, interconnections between the distinct pathways were found to be unique for each lineage, while one of such interconnections was essential for erythrocytes. Taken together, we here establish an HSPC differentiation model, suitable for metabolic toxicity screening and we assessed the unique combination of active metabolic pathways as well as their interconnections for each lineage, enabling the understanding of differing metabolic changes in myeloid hematopoiesis, caused by exogenous compounds.

## Results

### Expansion approach preserves restricted myeloid-lineage potential

As MS-based quantification of metabolites typically requires large amount of biological material, we applied an initial expansion step of CD34^+^ HSPC. The applied protocol leads to a total expansion of alive cells by 286±46-fold after 13 days (Figure 1B).^34^ However, only around 45% of total cells remain CD34^+^ positive after expansion, also primitive markers as CD135 and CD45RA largely remain negative while a large part of the population indicates CD90 positive (Figure 1A). Nevertheless, many typical lineage markers remain negative, only around 10% are CD235a positive (Supplemental Figure 1A).^35^ These results indicate that differentiation of primitive progenitors occurs during expansion, lack of most lineage markers, however, indicate a multi-lineage potential in the expanded population.

**Figure 1.**
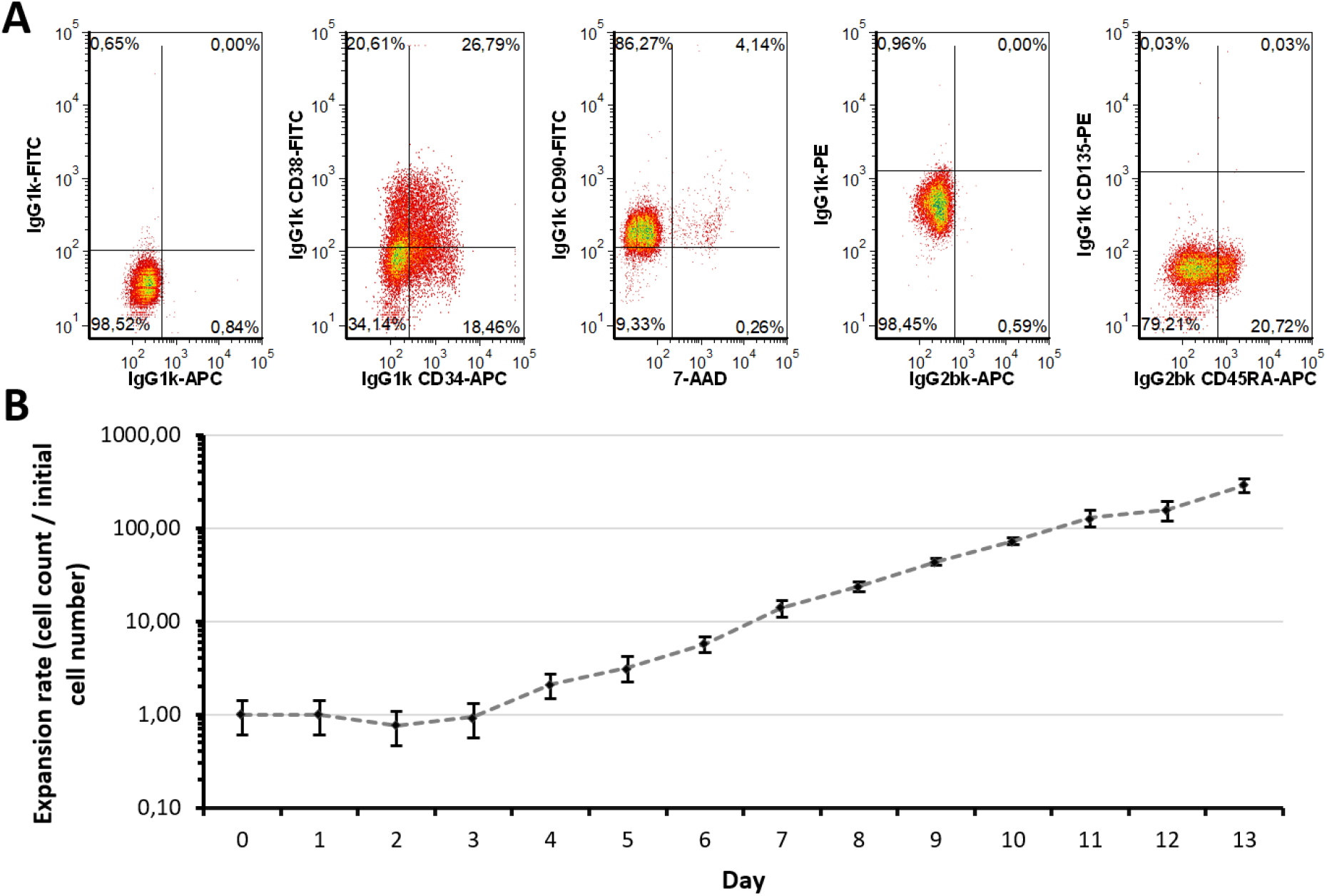
Characteristics of expanded CD34^+^ HSPC population. (A) Expression of HSPC surface markers in the progenitor population in comparison to corresponding isotype controls. Representative dot plots from three independent experiments (n=3) are shown. (B) Growth characteristics of CD34^+^ HSPC during expansion phase. Alive cell numbers were determined by trypan blue exclusion method, using a hemacytometer (n=3).

We, therefore, continued differentiation of the progenitor population into erythroid^36^, megakaryocytic^37^ and natural killer T cell lineages^38^. Erythroid differentiation could be confirmed (Figure S1b) while, lineage markers for megakaryocytic and natural killer T cell lineages remained negative (Figure S1c,d). Assessment of further lineage markers, however, revealed that CD1a^+^ dendritic cells (Figure S1c) and CD66b^+^ neutrophils (Figure S1d) were formed as major populations. This observation is also supported by transcriptomic data, as several of the most extensively expressed genes could be matched to the afore mentioned lineages (Table S1). As *HBB* and *TFRC* are both erythroid, *ELANE, S100-A8, PRTN3* and *AZU1* are neutrophil markers, *GPNMB* is expressed by antigen-presenting cells (APC), *MRC1* and *C1QC* by macrophages and DC, while *EPX* is expressed by eosinophils, the expression patterns indicate, that the main lineages formed are erythrocytes, DC as well as neutrophils.^39–43^ Furthermore, also gene ontology enrichment analysis of the generated populations support this assumption, as most significantly upregulated biological processes include erythrocyte differentiation and homeostasis for the erythroid population, as well as biological processes related to immune response for the other populations (Table 1).

**Table 1.**
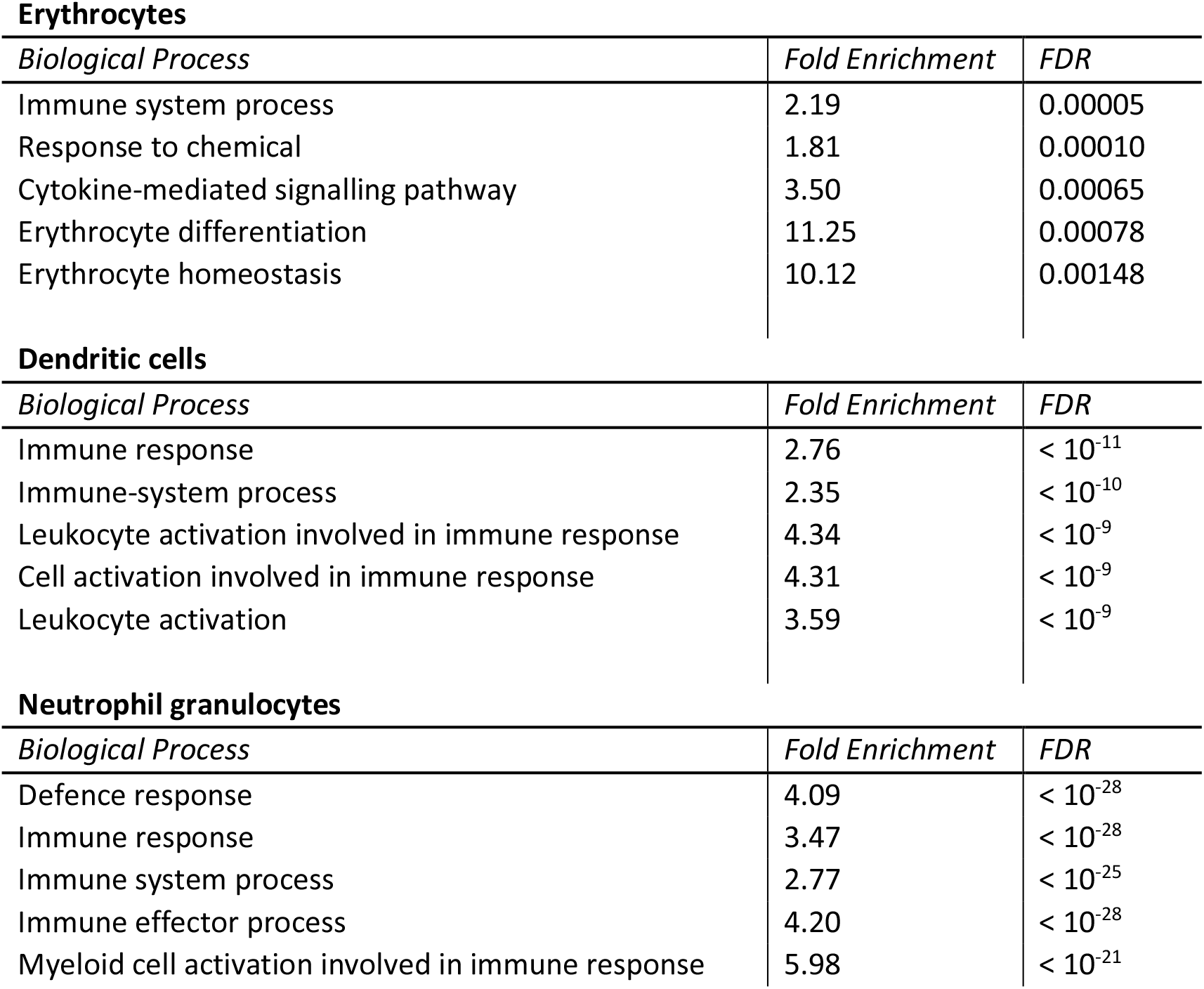
Gene Ontology Enrichment Analysis from the differentiated lineages. The five most significant enriched biological processes in each lineage are shown.

**Erythrocytes**
| *Biological Process* | *Fold Enrichment* | *FDR* |
| --- | --- | --- |
| Immune system process | 2.19 | 0.00005 |
| Response to chemical | 1.81 | 0.00010 |
| Cytokine-mediated signalling pathway | 3.50 | 0.00065 |
| Erythrocyte differentiation | 11.25 | 0.00078 |
| Erythrocyte homeostasis | 10.12 | 0.00148 |

**Dendritic cells**
| *Biological Process* | *Fold Enrichment* | *FDR* |
| --- | --- | --- |
| Immune response | 2.76 | $< 10^{-11}$ |
| Immune-system process | 2.35 | $< 10^{-10}$ |
| Leukocyte activation involved in immune response | 4.34 | $< 10^{-9}$ |
| Cell activation involved in immune response | 4.31 | $< 10^{-9}$ |
| Leukocyte activation | 3.59 | $< 10^{-9}$ |

**Neutrophil granulocytes**
| *Biological Process* | *Fold Enrichment* | *FDR* |
| --- | --- | --- |
| Defence response | 4.09 | $< 10^{-28}$ |
| Immune response | 3.47 | $< 10^{-28}$ |
| Immune system process | 2.77 | $< 10^{-25}$ |
| Immune effector process | 4.20 | $< 10^{-28}$ |
| Myeloid cell activation involved in immune response | 5.98 | $< 10^{-21}$ |

Taken together, these data demonstrate the multi-lineage potential of the generated progenitor population as it was possible to form erythroid, DC, neutrophil and eosinophil lineages. Since all populations reside in the myeloid branch of hematopoiesis, we assume that the progenitor population either consists of restricted myeloid progenitor cells or of a mixture of further differentiated progenitors. Indeed, transcriptomics data indicates that also other lineages are formed within each population, however, the main lineages formed are erythrocytes, DC as well as neutrophils, as verified via flow cytometry.

### Myeloid lineages display several unique and common metabolic patterns during differentiation accompanied by changes in pathway-related genes

Based on the successful identification of each main lineage formed, we performed global targeted metabolome profiling, testing for 514 metabolites in total. We totally identified 173 metabolites being altered from the progenitor population in the mature lineages (Figure 2A). In this set, 81 metabolites are significant in erythrocytes, 99 metabolites in DC and 73 in neutrophils (Supplemental Table 2-4). While several metabolic changes seem to be comparable between two or more populations, several unique changes are present (Figure 2A). By applying criteria for identification of unique and common changes, as stated in the methods section, we found 17 unique metabolites changed in erythrocytes (Figure 2A highlighted with red, Supplemental Table 5), 14 unique metabolites in DC (Figure 2A highlighted with green, Supplemental Table 6) and 46 unique metabolites in neutrophils (Figure 2A highlighted with blue, Supplemental Table 7). When these unique metabolic changes are directly compared in all three lineages, a cluster for each lineage is clearly visible (Figure 2B). Several metabolite changes in fact are found in common, with erythrocytes and DC displaying the largest overlap with 25 metabolites (Figure 2A highlighted with yellow, Supplemental Table 8). Common metabolites between the other combinations of populations, however, were rather rare (Figure 2A highlighted with cyan, pink and black, Supplemental Table 9-11).

**Figure 2.**
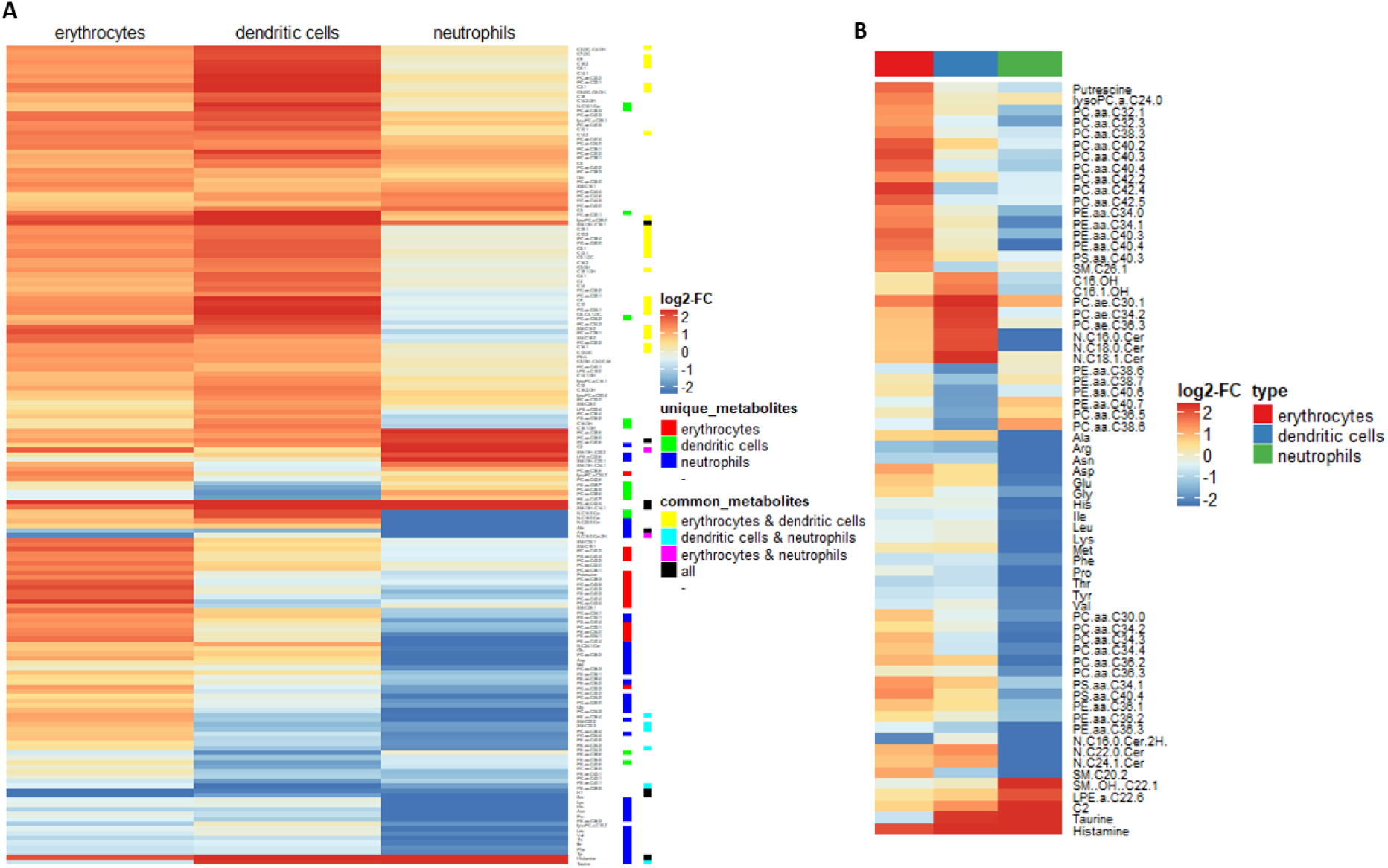
Biologically relevant metabolite changes in each differentiated lineage and identified unique metabolite patterns. (A) Biological relevant metabolite changes in each differentiated lineage. (B) Unique metabolite patterns in each differentiated population. The mean log_2_ fold change in comparison to the progenitor population of identified unique metabolites is displayed (n=3). Biologically relevant, unique and common metabolites were identified as stated in materials and methods section.

As we sought to identify the underlying changes on metabolic pathway levels, we analysed the expression of enzymes, related to the relevant metabolites. Whole pathway activity is often regulated by a small number of key enzymes (e.g. hexokinase, phosphofructokinase and pyruvate kinase in glycolysis), therefore, whole pathway activity can often be derived from the expression of these in combination with metabolite concentrations.^44^ Furthermore, as enzyme activity is tightly controlled via several mechanisms, including expression, we omitted FDR-correction of the p value in order to avoid an increase of the type II error (false negatives).^45^ By using this approach, we found several transcriptional changes, related to the corresponding metabolic pathways (Table 2).

**Table 1.** Transcriptional changes in pathway related genes in each population. Given is gene name, the corresponding metabolic pathway and the log_2_ fold change, compared to the progenitor population. Changes with a corresponding p-value < 0.05 were considered significant.

| Gene | Metabolic pathway | Erythrocytes | Dendritic cells | Neutrophils |
| --- | --- | --- | --- | --- |
| *SLC2A1* | Glycolysis | 1.69 | n.s. | n.s. |
| *SLC2A5* |  | n.s. | n.s. | 1.94 |
| *SLC2A6* |  | n.s. | n.s. | 2.28 |
| *HK3* |  | 1.77 | 2.65 | 3.76 |
| *MINPP1* |  | 0.96 | -1.47 | n.s. |
| *PKLR* |  | 4.42 | n.s. | n.s. |
| *FBP1* |  | n.s. | 4.44 | 5.36 |
| *OAT* | Polyamine synthesis | 0.95 | n.s. | n.s. |
| *ODC* |  | n.s. | -1.89 | -1.47 |
| *SAT1* |  | n.s. | 1.28 | 1.16 |
| *MYC* |  | 1.19 | n.s. | n.s. |
| *ME1* | NADPH production | n.s. | 2.88 | n.s. |
| *IDH1* |  | n.s. | 1.97 | n.s. |
| *CPT1a* | FA oxidation | n.s. | n.s. | -1.25 |
| *FASN* | FA biosynthesis | 1.10 | n.s. | n.s. |
| *ACSL1* |  | n.s. | 1.14 | n.s. |
| *ELOVL6* | FA elongation | 1.41 | n.s. | n.s. |
| *THEM5* |  | n.s. | n.s. | Inf |
| *SCD* | Biosynthesis of unsaturated FA | 1.13 | 1.36 | n.s. |
| *LIPA* | Lipolysis | n.s. | 2.86 | 2.75 |
| *PLD3* | Glycero-phospholipid metabolism | 0.96 | 2.54 | 2.33 |
| *PLBD1* |  | 1.84 | n.s. | 3.118 |
| *PLA2G15* |  | n.s. | 1.24 | n.s. |
| *PLA2G16* |  | n.s. | n.s. | 2.73 |
| *LPCAT3* |  | 1.09 | n.s. | n.s. |
| *PISD* |  | 1.03 | n.s. | n.s. |
| *PHOSPHO1* |  | 3.91 | n.s. | n.s. |
| *PPAP2B* |  | -2.28 | 1.73 | 2.54 |
| *LPIN1* |  | n.s. | n.s. | -Inf |
| *PCYT1B* |  | n.s. | -2.02 | -2.40 |
| *PLA2G7* | Ether lipid metabolism | 1.41 | 4.18 | 3.34 |
| *SPTSSB* | Sphingolipid metabolism | 3.11 | 5.07 | n.s. |
| *ACER3* |  | n.s. | 1.14 | n.s. |
| *SMPD3* |  | Inf | n.s. | Inf |

### Myeloid lineages own a higher hexose consumption with different fate

As the energy metabolism seems to play a crucial role during HSPC maintenance and differentiation, we focused on glycolysis, FAO and glutaminolysis as these display the three major pathways for energy transformation.^46^ Hexoses were found to be strongly reduced in all lineages (Supplemental Table 12), also expression of corresponding transmembrane transporters (*SLC2A1, SLC2A5* and *SLC2A6*) was either upregulated or unchanged (Table 2) indicating that lowered hexoses are related to raised catabolic reactions. This interpretation is further supported by the increased expression of *HK3* in all lineages. However, expression of *PKLR* was upregulated in erythrocytes, while *FBP1* was upregulated in DC and neutrophils. As *PKLR* catalyse the formation from phosphoenolpyruvate to pyruvate, it can be concluded that erythrocytes own higher consumption of hexoses, subsequently used for the Krebs cycle. It has, however, been shown that nucleotide synthesis from glucose-6-phosphate (G6P) via the PPP is vital for erythroid differentiation.^29^ However, we could not find any difference in the expression of enzymes related to the PPP, indicating that erythrocytes use G6P under native conditions for further catabolic breakdown via glycolysis, as well as for nucleotide biosynthesis via PPP. *FBP1* catalyses the back conversion of fructose 1,6-bisphosphate to fructose 6-phosphate (F6P) and acts as the rate-limiting enzyme in gluconeogenesis. As F6P readily can be converted to G6P, we assume that both lineages primary use glucose for nucleotide biosynthesis and ROS production from NADPH via PPP, which is also supported by findings of other groups.^47,48^

### Combined fatty acid generation and respective fate is unique for each myeloid lineage

Several acylcarnitines were raised in erythrocytes and DC, 18 of which could be defined as change common to both lineages (Supplemental Table 3,4 and 9). Only three acylcarnitines, however, were increased in the neutrophil lineage, with acetylcarnitine concentration being uniquely affected (Supplemental Table 5 and 8). Acylcarnitines serve as carnitine shuttle for fatty acids through the mitochondrial membranes, while even-chain acylcarnitines and hydroxylated species result from β-oxidation, odd-chain species from α-oxidation and dicarboxylated species from ω-oxidation.^49–52^ It is well known that acylcarnitines can also play a role in maintaining free CoA availability, whilst also reflecting corresponding acyl-CoA concentrations within the cell.^50^ As activity of FAO was demonstrated in HSC and during neutrophil differentiation, it is reasonable to assume that both possess enhanced FAO.^27,46^ We therefore assume, that raised acylcarnitines indicate a lower FAO, which is also supported by the fact that species, resulting from α- and ω-oxidation, are raised as well. As α- and ω-oxidation only take place at substantial amounts if β-oxidation is disturbed, this further indicates reduced FAO.^53,54^

Conversely, we found expression of *CPT1a* lowered in the neutrophil lineage (Table 2). As transesterification by *CPT1a* is one of the rate-limiting steps in FAO, this decreased expression seems contradictory to our assumptions. However, usage of etomoxir as irreversible inhibitor of *CPT1* only affected neutrophil differentiation (Figure 3A,D and 4B,C), confirming our hypothesis. Surprisingly, addition of etomoxir during neutrophil differentiation led to higher expression of *ELANE* (Figure 3D). As etomoxir effectively reduced the proportion of CD11b^+^ cells during neutrophil differentiation, our results seem rather conflicting.^27^ However, *ELANE* and CD11b are both expressed at different stages during neutrophil maturation, while *ELANE* being expressed in promyelocytes and myelocytes; CD11b expression starts in metamyelocytes.^55^ Furthermore, expression of *ELANE* is inversely correlated to maturation of neutrophils.^56^ Therefore, we conclude, that higher expression of *ELANE* in etomoxir treated populations indicates an enrichment of promyelocytes and myelocytes. In line with this assumption, the expression of *S100A8*, correlating with neutrophil maturation, was lowered by increasing amounts of etomoxir (Figure 3E).^56^ Thus, FAO seems to be essential for neutrophil maturation, whereas, formation of committed progenitors seems to be independent of FAO. This interpretation, however, awaits further confirmation by future studies.

**Figure 3.**
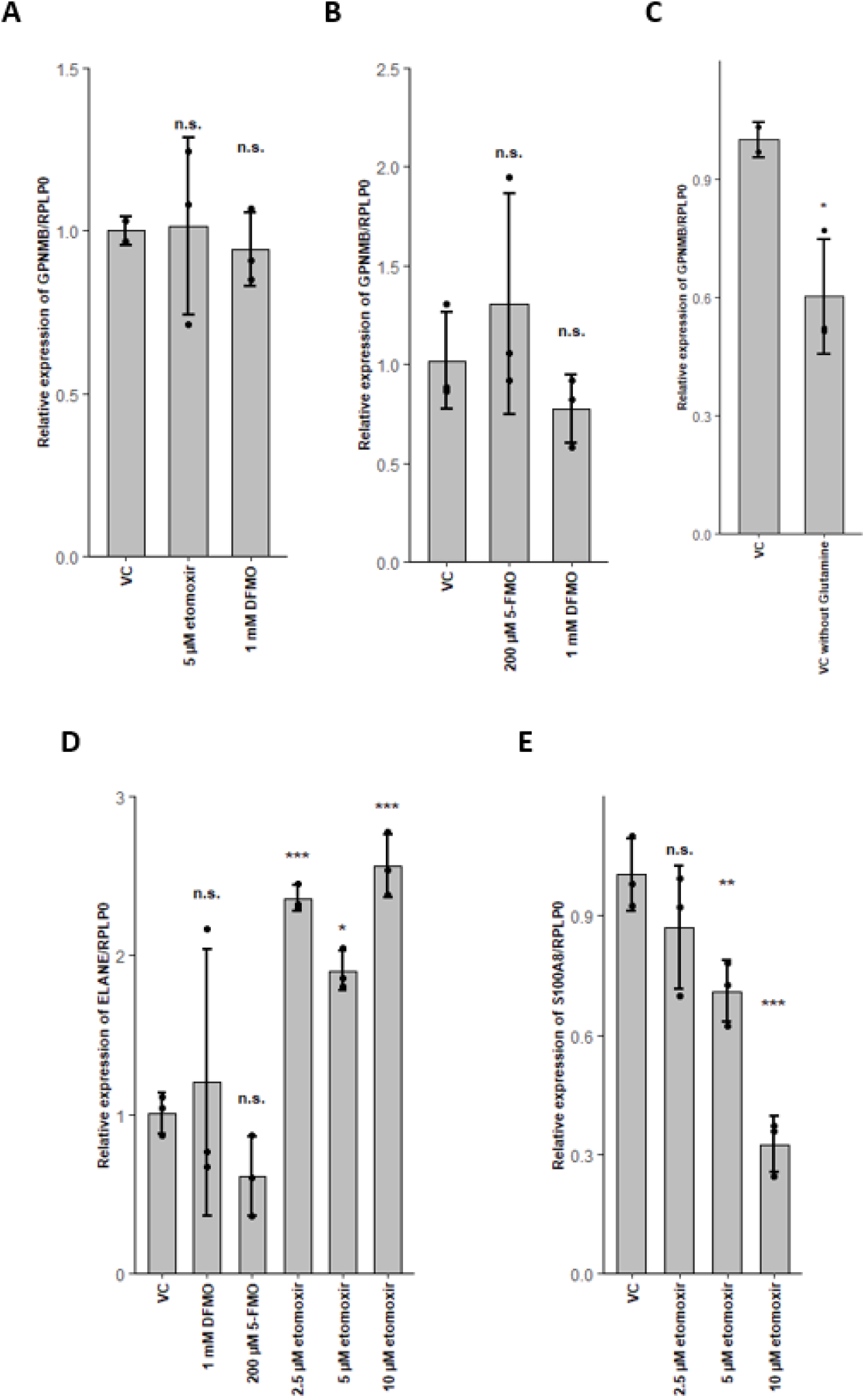
Glutaminolysis is essential during dendritic cell differentiation, while fatty acid oxidation is essential for neutrophil maturation. (A) Effect of etomoxir or difluormethyl-ornithine (DFMO) in presence of glutamine on GPNMB/RPLP0 expression of dendritic cells. (B) Effect of 5-fluoromethylornithine (5-FMO) or DFMO in absence of glutamine on GPNMB/RPLP0 expression of dendritic cells. (C) Effect of glutamine on GPNMB/RPLP0 expression of dendritic cells. (D) Effect of 5-FMO, DFMO or etomoxir in presence of glutamine on ELANE/RPLP0 expression of neutrophils. (E) Effect of etomoxir in presence of glutamine on S100A8/RPLP0 expression of neutrophils. Data are represented as mean ± SD (n = 3). Relative gene expression was calculated as stated in the methods section. Significant changes were assessed by one-way ANOVA (n.s. not significant; *p<0.05; ***p<0.001).

Interestingly, C16-OH and C16:1-OH acylcarnitines were uniquely raised in DC, but not in erythrocytes (Figure 2B, Supplemental Table 8). In fact, intracellular fatty acids (FA) and corresponding acyl-CoA species can result from catabolic reactions via lipolysis, as well as from anabolic reactions via FA synthesis. We found expression of *FASN* and *ELOVL6* uniquely stimulated in the erythrocyte lineage, while expression of *LIPA* was enhanced in DC and neutrophils. Furthermore, we found *ACSL1* elevated in DC and *THEM5* in neutrophils, also *SCD* was elevated in erythrocytes and DC.

Summarized, these results indicate that erythrocytes perform higher FA synthesis, while both DC and neutrophils perform higher lipolysis of cholesteryl esters (CE) and triglycerides (TG). FAs are further catabolised via FAO in neutrophils, while erythrocytes and DC seem to utilize FAs for anabolic processes. Erythrocytes, however, preferably elongate C16:0 and C16:1 FAs (Figure 5), which leads to diminished concentrations of these FAs, which is, in turn, reflected by corresponding acyl-CoAs and thus acylcarnitines.

### Glutaminolysis crosstalk with FAO and its interconnection to polyamine synthesis in erythrocytes

In context to glutamine metabolism, we found uniquely lowered levels of the products Glu, Asn, Asp, Ala and Pro in neutrophils (Figure 2B and 5). As it is known that glutaminolysis is critical to erythroid differentiation, we assume that both, erythrocytes and DC perform glutaminolysis.^29^ This assumption is further supported by reduced *GPNMB* expression in DC upon glutamine depletion (Figure 3C). In erythrocytes, however, glutamine depletion only reduced the cell expansion rate, *HBB* expression remained unaffected (Figure 4H,I). Glutaminolysis, therefore, seems to be crucial for DC differentiation, in erythrocytes, it, however, appears to be beneficial for cell expansion but not for maturation. In neutrophils, however, we assume that high ATP production by FAO concomitantly leads to reduced free phosphate levels, which in turn lead to lowered glutaminase activity.^57^

**Figure 4.**
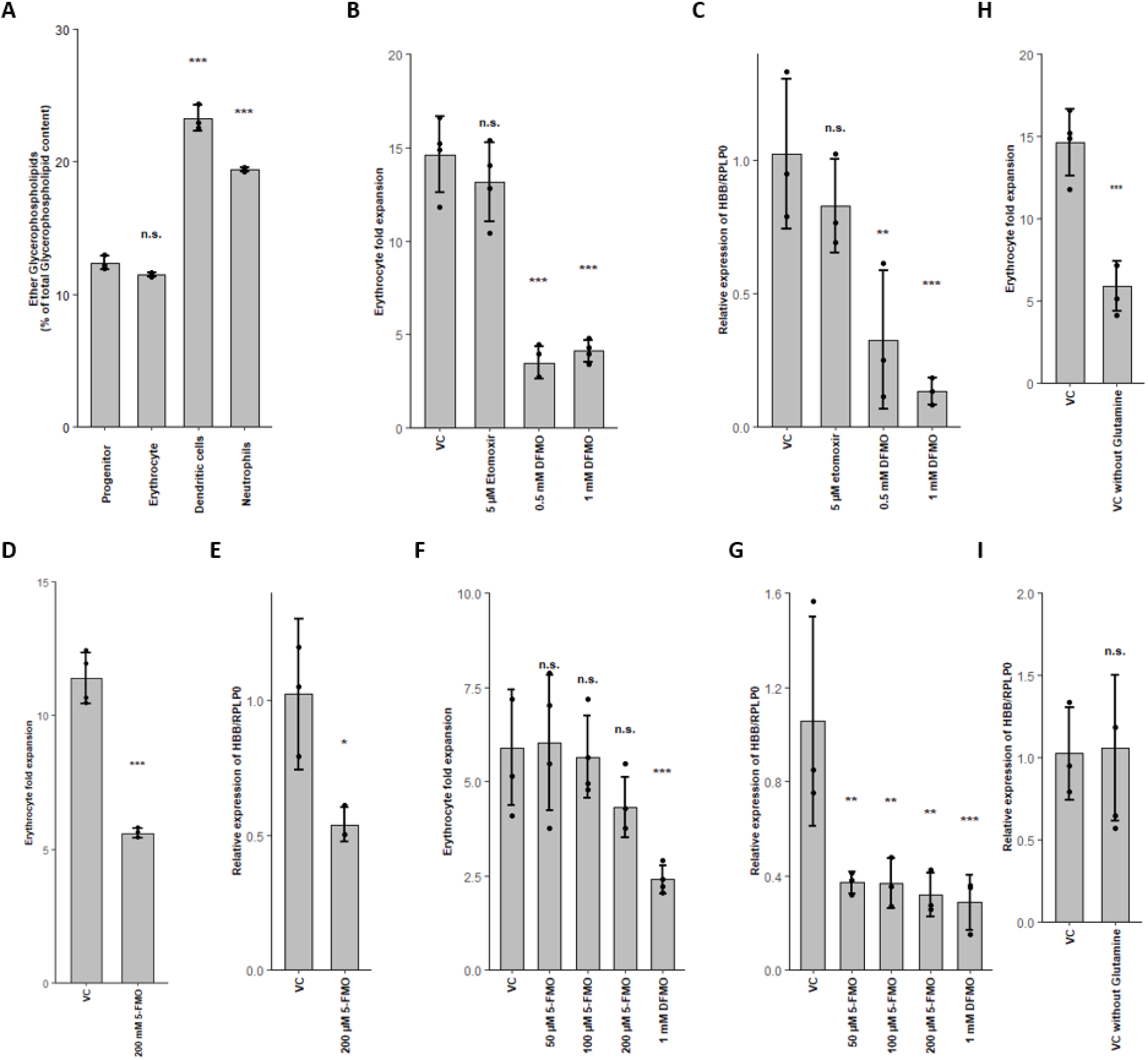
Polyamine synthesis and glutamine metabolism are both essential for erythropoiesis and interconnected via *OAT*. (A) Percentual content of ether lipids. (B and C) Effect of etomoxir or difluormethyl-ornithine (DFMO) in presence of glutamine on erythroid expansion and relative HBB/RPLP0 expression. (D and E) Effect of 5-fluoromethylornithine (5-FMO) in presence of glutamine on erythroid expansion and relative HBB/RPLP0 expression. (F and G) Effect of 5-FMO or DFMO in absence of glutamine on erythroid expansion and relative HBB/RPLP0 expression.(H and I) Impact of glutamine on erythroid cell expansion and relative HBB/RPLP0 expression. Data are represented as mean ± SD (n = 3 for ether lipid content and gene expression, n = 4 for cell expansion). Percentual content of lipids, expansion rates and relative gene expression was calculated as stated in the methods section. Significant changes were assessed by one-way ANOVA (n.s. not significant; *p<0.05; **p<0.01; ***p<0.001).

Interestingly, we found *ME1* and *IDH1* raised in DC, both resulting in NADPH generation from Krebs cycle intermediates (Table 2).^19^ Cross-regulation between glycolysis and FAO via NADPH and acetyl-CoA is well known, furthermore, pyruvate is generated via *ME1*.^58^ We therefore assume, that in DC, FAO is mainly negatively regulated by glutaminolysis and not glycolysis (Figure 5).

**Figure 5.**
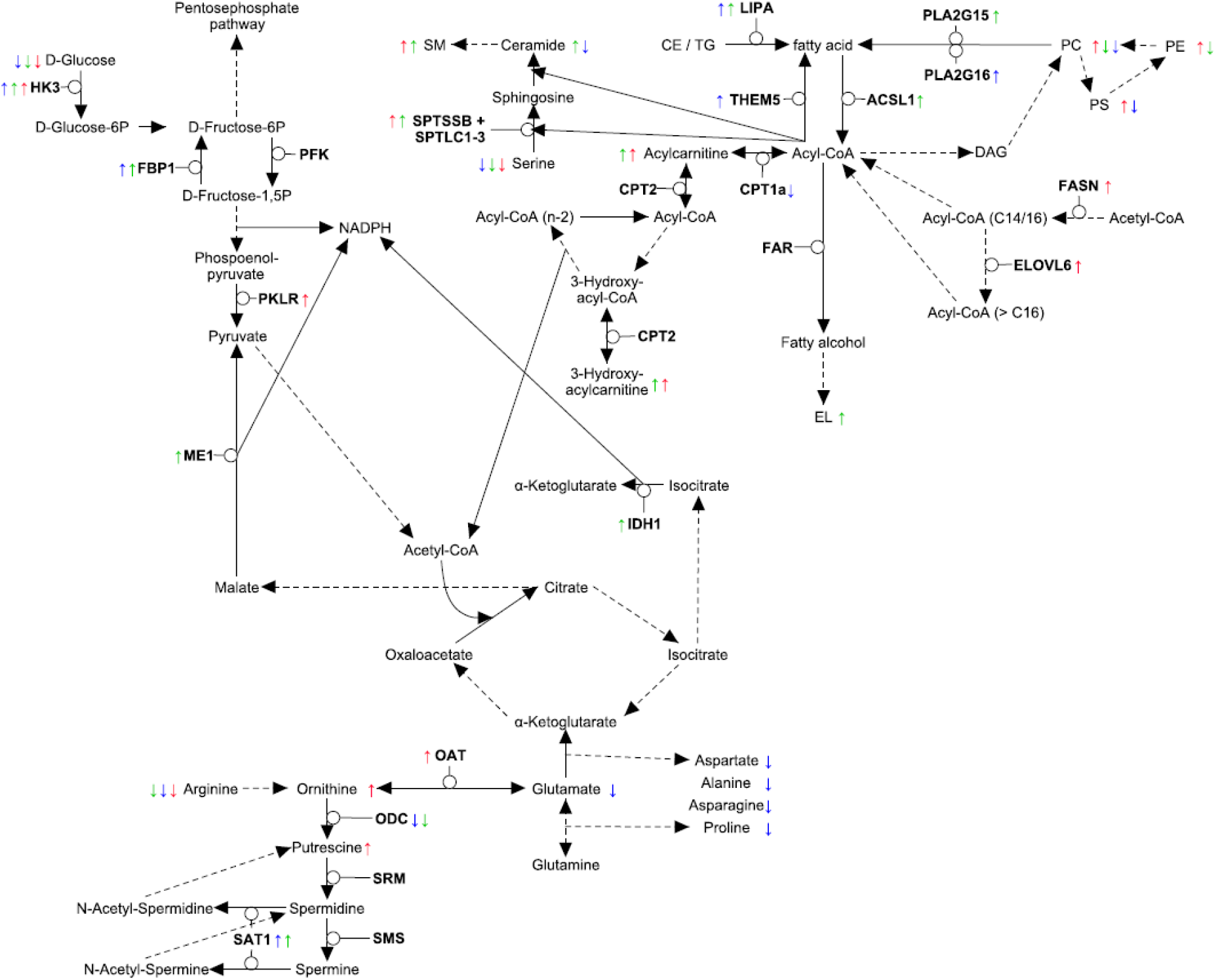
Involvement of unique metabolites in interconnected metabolic pathways. Model depicting the interconnection of key reactions of glycolysis, Krebs cycle, glutaminolysis, FAO and lipid metabolism. Differences found are highlighted for neutrophils (blue), dendritic cells (green) and erythrocytes (red).

Furthermore, we found putrescine uniquely raised in erythroid differentiation (Figure 2B, Supplemental Table 6), while downstream metabolites spermidine and spermine remained unaffected. Also, Orn was elevated in erythrocytes whilst not fulfilling the criteria as unique metabolite, while Arg was lowered in all lineages (Figure 2B, Supplemental Table 3,8 and 12). We further found *ODC* expression reduced in DC and neutrophils, while *SAT1* was raised (Table 2). Polyamine levels in fact have been reported to be relatively high in erythrocytes, confirming our observations.^59^ This pattern indicate that anabolism of polyamines is stable during erythroid differentiation, resulting in raised putrescine levels, whilst being diminished during DC and neutrophil differentiation. Several reason may account for this: It is well known, that polyamines are important in regulating gene expression, as they are mostly bound to RNA and are essential for cell cycle progression.^33^ As such they are found at higher concentrations in high proliferating cells. Further, polyamines were found to inhibit transbilayer movement of phospholipids and reduce lipoperoxidation in erythrocytes. ^60–62^ The importance of polyamines during erythropoiesis was also previously demonstrated by other research groups.^59,63^ In line with this, inhibition of polyamine synthesis by blockage of *ODC* via difluormethyl-ornithine (DFMO) reduced erythroid expansion rates and *HBB* expression (Figure 4B,C). DC and neutrophils, in contrast, were not affected by *ODC* inhibition (Figure 3A,D). Therefore, polyamine synthesis seems solely crucial for erythropoiesis, at least in the myeloid branch of hematopoiesis. Until now, however, it is not exactly clear which of the stated functions of polyamines is crucial for erythropoiesis; we assume that all of them might be crucial for different stages during erythropoiesis.

Interestingly, we found the expression of *OAT* solely increased during erythropoiesis (Table 2). *OAT* catalyses the generation of Glu and glutamate-5-semialdehyde from Orn and aKG and vice versa.^64^ Under most conditions, *OAT* uses Orn for anabolism of Glu and, therefore, belongs to the “glutamate crossway”. Under certain conditions, however, *OAT* can switch to Orn synthesis. As Glu remained unchanged and Orn was raised during erythropoiesis, we, however, assume that the favoured reaction is Glu anabolism. In line with this, it has been shown that erythropoiesis is sustained during depletion of extracellular Gln, but is dependent on the intracellular synthesis from Glu under these conditions.^29^

In order to evaluate the influence of *OAT* on erythropoiesis, as well as the direction of the conversion, we used 5-fluoromethylornithine (5-FMO), a specific inhibitor of *OAT*.^65^ Surprisingly, addition of 5-FMO lead to an reduced expression of *HBB* independent of Gln (Figure 4E,G), whilst expansion rate was only affected in presence of Gln (Figure 4D,F). We, therefore, propose, that *OAT* works in both directions, depending on the conditions; in presence of Gln *OAT* catalyses the conversion of Glu to Orn, subsequently fuelling polyamine synthesis, whilst under conditions with low Gln, *OAT* prefers the conversion of Orn to Glu, leading to constant levels of Gln for nucleotide biosynthesis (Figure 5).^29^ The homeostasis of polyamine synthesis and Gln levels is evident, as inhibition of *ODC* by DFMO leads, independent of Gln levels, to diminished *HBB* expression (Figure 4C,G). Additionally, inhibition of *OAT* in absence of Gln also reduced *HBB* expression to similar levels as DFMO, indicating that polyamine synthesis alone is not sufficient for erythropoiesis (Figure 4G). Importantly, we found that inhibition of *OAT* or *ODC* in absence of Gln did not influence *GPNMB* expression of DC (Figure 3B), also expression of *ELANE* by neutrophils remained unaffected by 5-FMO (Figure 3D). Therefore, *OAT* mediated homeostasis of polyamine synthesis and Gln seems to be only crucial during erythropoiesis.

### PLA2G15, PLA2G16 and fatty acid levels are three major regulators of unique lipidomic changes

As several uniquely changed metabolites were glycerophospholipids (GPL), we also focused on GLP metabolism. GLP need to be further subdivided in acyl-acyl species (acyl glycerophopholipids, aGPL) and acyl-alkyl species (ether lipids, EL), as former are involved into glycerophospholipid metabolism and the latter are involved into ether lipid metabolism. Interestingly, the different GPL classes (e.g. phosphatidylcholine (PC), phosphatidylethanolamine (PE) and phosphatidylserine (PS)) followed the same trend in all lineages (not shown). However, erythrocytes display increased aGPL levels, DC show lowered aGPL levels and higher EL levels while neutrophils exhibit reduced aGPL levels, in comparison to the progenitor population (Figure 2 A,B, Supplemental Table 5-7). Furthermore, percentual content of EL was elevated in DC and neutrophils (Figure 4A), as also reported previously.^66^

We actually found several changes in the expression of enzymes, related to glycerophospholipid and ether lipid metabolism (Table 2), however, only *PLA2G15*, *PLA2G16, LPCAT3, PISD, PHOSPHO1* and LPIN1 were affected distinctively in the corresponding populations. Furthermore, activity of *PLBD1, PLD3* and *PISD* has been reported as negligible, which is also reflected by the corresponding metabolic changes.^67–70^ As we found enhanced *de-novo* FA synthesis and elongation in erythrocytes, as well as stimulated lipolysis in DC and neutrophils, the unique metabolite patterns apparently partially reflect available FA levels, as longer aGPL’s are raised in erythrocytes. In DC and neutrophils, aGPL’s seem to be degraded by *PLA2G15* and *PLA2G16*.^71,72^ The available FA levels are further reflected by EL’s, as *FAR1/2*, the major regulator of EL biosynthesis, uses C16 and C18 acyl-CoA species as substrates.^73,74^ Since C16 FA are further elongated in erythrocytes, this results in limited EL synthesis due to limited substrate availability and thus unchanged EL levels. Further, in neutrophils, *PLA2G16* reduces EL biosynthesis by inducing peroxisomal dysfunction, which results in lack of finding uniquely raised EL, despite their higher percentual content.^72^

In line with this, we found higher levels of C16, C18 and C18:1 ceramide in DC (Figure 2B). Furthermore, expression of *SPTSSB*, well-known to stimulate serine palmitoyltransferase *(SPT)* activity, was raised in DC and erythrocytes.^75^ As *SPT*uses C18-CoA substrates, we conclude that *de-novo* ceramide synthesis is enhanced in DC due to higher availability of sphingosine, resulting from higher C18-CoA availability.

Taken together, our data strongly indicates that parts of the cellular lipidome analysed here are all regulated by FA levels, in combination with expression of *PLA2G15* or *PLA2G16*, resulting in the described unique metabolic signatures.

## Discussion

Cellular metabolism is an essential feature of living organisms, various metabolites are involved in specialized cellular tasks, ultimately defining the function of each cell. In context to hematopoietic stem cells, it is well appreciated that activity of metabolic pathways not only regulate quiescence, but also regulate commitment.^46,76^ Inhibition of FAO, for example, leads to symmetric commitment of HSCs, however, neutrophil differentiation is also disturbed by inhibition of FAO. ^26,27^ The regulatory role of FAO, therefore, remains controversial to some extent. Furthermore, several studies have shown the importance of various metabolic pathways on differentiation of several lineages.^29,30,59,77^ However, the lineage-specific activity of these metabolic pathways remains vague, a direct metabolic comparison of distinct lineages is still missing.

The total cellular metabolism plays a fundamental role in the cellular response towards different stimuli. In context to hematopoiesis, several different compounds are known to exert distinct effects on the different blood cell populations.^9,24,78–80^ The actual mode of action behind this lineage specific hematotoxicity, however, often remains inconclusive. As involvement of the total cellular metabolism in such lineage specific effects is quite likely, fundamental knowledge of the different active and inactive metabolic pathways is essential to understand such lineage specific effects.

We, therefore, established an HSPC differentiation model, suitable for omics-based screening approaches, and performed metabolic and transcriptomic comparison of three myeloid lineages, regarding to the initial progenitor population. Generated populations in fact were not purified further, potentially leading to overlapping metabolic signatures. However, by identifying distinctively altered metabolites, we found several lineage-related active metabolic pathways, some of which have also been identified previously (e.g. polyamine synthesis in erythrocytes^59^ and FAO in neutrophils^27^). Presence of other lineages is even of advantage from a toxicological point of view, as toxic effects on these can also be identified simultaneously. The corresponding mode of action, then may be identified within the dominant lineage, using omics techniques. We found assumptions made from metabolomic and transcriptomic data in good agreement with results of verifying inhibitor experiments, indicating the validity of the here described approach.

We could further demonstrate that not only FAO, but also the combination of fatty acid generation and the fate is unique for each lineage. Also, inhibition of FAO did impair neutrophil maturation, but not formation of committed progenitors, supplementing current knowledge about FAO dependence of HSC commitment. Of note, our data also suggest that glycolysis and glutaminolysis have substantial impact on FAO, leading to differential usage of fatty acids in each lineage. We could further demonstrate a unique and essential connection of glutamine metabolism to polyamine synthesis in erythrocytes and identified the key players in lipid remodelling during myeloid lineage commitment, namely acyl-CoA availability, *PLA2G15* and *PLA2G16*.

Our results thus connect several crucial active metabolic pathways identified in single myeloid lineages, while also displaying their unique combination for each lineage, as well as their inactivity in the other lineages. Direct usability for hematotoxicity studies is evident, as several xenobiotic compounds are known to selectively interact with specific metabolic pathways.^81–83^ Likewise, elucidation of effects by endogenous compounds on hematopoiesis can also be achieved by usage of the here presented model, which may ultimately result in new therapy approaches. Furthermore, our results enable improved interpretation of metabolic alterations, observed by different compounds.^84–86^

Taken together, our study adds significant knowledge in the understanding of the differences in the metabolism of different myeloid lineages and enables transferability of results obtained with such model towards other tissues. It will aid future studies in understanding the differential impact of environmental or xenobiotic compounds on hematopoiesis and adds essential knowledge for identifying phenotype-modulating metabolites.

## Materials and methods

### Reagents and materials

Human CB CD34^+^ cells were purchased from STEMCELL Technologies GmbH (Cologne, Germany). StemPro-34 SFM, IMDM and valproic acid were from Fisher Scientific (Wien, Austria). Cytokines were purchased from GeneScript (Piscataway, NJ). BML-210 and stemregenin 1 (SR-1) were from ApexBio Technology LCC (Houston, TX). L-Glutamine was from Carl Roth GmbH+Co.KG (Karlsruhe, Germany). Etomoxir was from VWR International GmbH (Bruchsal, Germany). Elfornithine (DFMO) and Human male AB serum was from Sigma-Aldrich Chemie GmbH (Munich, Germany). 5-Fluoromethylornithine (5-FMO) was from Chemspace Europe (Riga, Latvia). All antibodies were purchased from Beckton Dickinson GmbH (Heidelberg, Germany).

### Flow cytometry analyses and antibodies

All flow cytometry data were acquired on a CyFlow Cube 8 flow cytometer (Sysmex) and analysed using FCS Express software (De Novo Software). Cells were washed twice with PBS and blocked with 10% human male AB serum for 15 min at 4°C. The following are the anti-human antibodies used: CD34-allophycocyanin (APC), CD38-fluorescein isothiocyanate (FITC), CD45RA-APC, CD135-phycoerythin (PE), CD71-FITC, CD235a-APC, CD3-APC, CD94-FITC, CD41-APC, CD42a-FITC, CD90-FITC, CD66b-FITC, CD14-Pacific Blue (Pac Blue) and CD1a-APC, FITC-, PE-, APC- or Pac Blue-conjugated isotype-matched antibodies served as controls.

### Human CD34^+^ cell culture

CD34^+^ cells were thawed according to the vendors protocol and expanded in Stempro-34 SFM media containing 2 mM L-glutamine, 100 ng/ml SCF, 100 ng/ml Flt-3L, 50 ng/ml IL-6, 40 ng/ml TPO, 1 μM SR-1, 0.1 μM BML-210 and 0.2 mM valproic acid for 13 days.^34^ Until day 8, each second day 5 ml fresh media were added to the culture. From day 9 to 13, cells were diluted 1:1 each day by addition of fresh media. Erythroid differentiation was performed in Stempro-34 SFM media containing 2 mM L-glutamine, 100 ng/ml SCF, 100 ng/ml Flt-3L, 50 ng/ml EPO and 20 ng/ml IL-3 for a total of 6 days.^36^ Cells were diluted 2:3 on day 4 by addition of fresh media. DC differentiation was done in Stempro-34 SFM media containing 2 mM L-glutamine, 10 ng/ml SCF, 10 ng/ml TPO and 10 ng/ml IL-3 for a total of 11 days. Neutrophil differentiation was done in IMDM containing 10% FBS, 50 ng/ml SCF and 50 ng/ml IL-15 for a total of 10 days. Dendritic cells and neutrophils were diluted 2:3 on day 3 and 7 by addition of fresh media. Different inhibitors were added as indicated at the beginning of each differentiation and held constant during the experiment, the added volume did not exceed 0.1% (v/v).

### Total RNA isolation and RNA-Seq

Total RNA isolation was performed using the NucleoSpin RNA XS Kit (Macherey-Nagel GmbH &Co.KG). Purification of poly-A containing mRNA molecules followed by mRNA fragmentation, random primed cDNA synthesis and single read 50 bp sequencing, as well as, transcriptome alignment and determination of differential expression levels were done by Eurofins GATC Biotech GmbH (Constance, Germany).^87^

### Quantitative PCR primer

HBB: fw: 5’-CTCGCTTTCTTGCTGTCCA-3’, rv: 5’-CAAGGCCCTTCATAATATCCCC-3’; ELANE: fw: 5’-CTGCGTGGCGAATGTAAACG-3’, rv: 5’-CGTTGAGCAAGTTTACGGGG-3’; GPNMB: fw: 5’-CTGATCTCCGTTGGCTGCTT-3’, rv: 5’-CTGACCACATTCCCAGGACT-3’; S100A8: fw: 5’-GATAAAGATGGGCGTGGCAG-3’, rv: 5’-TGCAGGTACATGTCCAGGG-3’; RPLP0: fw: 5’-TGGCAATCCCTGACGCACCG-3’; rv: 5’-TGCCCATCAGCACCACAGCC-3’.

### LC/MS-based metabolomics

Metabolite measurement was performed using a 4000 QTRAP mass spectrometer (ABI Sciex), connected to a NexteraXR HPLC (Shimadzu). Cell pellets were washed twice with ice-cold PBS and the pellets were resuspended at 1×10^7^ cells/ml in −20°C cold ethanol containing 15% (v/v) 10 mM K_2_PO_4_, pH 7.5, sonicated for 3 min and frozen in liquid nitrogen for 30 seconds. The sonication-freeze cycle was repeated twice, afterwards, samples were centrifuged at 31,500 rcf at 2°C for 5 min and the supernatant was stored at −80°C until measurement. The metabolome analyses were carried out using the AbsoluteIDQ p180 Kit (Biocrates Life Science AG), according to the manufacturer’s instructions. Analysis of additional lipids, covering 162 glycerophospholipids, 33 sphingomyelins and 131 ceramides was done by Biocrates Life Sciences AG (Innsbruck, Austria).

### Bioinformatic analysis

Statistical analysis of metabolomics data using an empirical Bayes approach was done by using the statistical software R^88^ and the bioconductor package limma^89^. Absolute log_2_ fold changes (logFC) above 1, compared to the progenitor population, with a corresponding FDR-adjusted p value below 0.05 where considered relevant. From these, unique metabolites were defined by a ΔlogFC > 1 to both other lineages. As ΔlogFC, we defined the absolute difference in the logFCs from each lineage compared to the progenitor population. Common metabolites of two lineages were defined as owning a ΔlogFC < 1, whilst both owning a ΔlogFC compared to third lineage > 1, from the relevant set of metabolites. Common metabolites of all lineages were defined as Δ logFC < 1 between all lineages from the relevant set of metabolites. Gene Ontology enrichment analysis was done by using the PANTHER Software (http://pantherdb.org).^90^ Significant changes were assessed by a one-way ANOVA, where a p value smaller than 0.05 was considered as significant. For metabolic pathway visualization, using KEGG pathways as template, the software PathViso^91,92^ was used.

### Data availability

RNA sequencing datasets generated are available at GEO under the accession number GSE129993. A list of identified significant altered metabolites can be found in the supplementary materials.

## Acknowledgments

This work was supported by a scholarship for Lars Kaiser from the Federal Ministry of Science, Research and Art of Baden-Württemberg, support by Steinbeis Center for Personalized Medicine (StZ1789) is gratefully acknowledged. MJ thanks the German Research Foundation (DFG) under Germany’s Excellence Strategy (CIBSS – EXC-2189 – Project ID 390939984) for support.

## Author Contributions

H.P.D., F.W. and M.J. were involved in study design. L.K. performed HSPC expansion and differentiation, inhibitor experiments, flow cytometry, metabolite and RNA extraction and mass spectrometric measurements. H.W. supported mass spectrometric measurements. I.Q. and M.B. performed primer design and quantitative RT-PCR. L.K. and M. K. performed bioinformatic analysis. L.K., H.P.D. and M.J. wrote the paper, all authors commented on the manuscript.

## Supplemental Information titles and legends

**Figure S1**. Expression of typical lineage markers in the different populations.

(A) Expression of typical lineage selective surface markers in the progenitor population in comparison to corresponding isotype controls, determined by flow cytometry. Representative dot plots from three independent experiments (n=3) are shown.

(B) Expression of erythroid (CD71 and CD235a) lineage surface markers in the erythrocyte population in comparison to corresponding isotype controls, determined by flow cytometry. Representative dot plots from three independent experiments (n=3) are shown.

(C) Expression of megakaryocytic (CD41a, CD42a), monocytic (CD14) and dendritic cell / Langerhans cell (CD1a) lineage surface markers in the dendritic cell population in comparison to corresponding isotype controls, determined by flow cytometry. Representative dot plots from three independent experiments (n=3) are shown.

(D) Expression of natural killer T cell (CD3, CD94), monocytic (CD14) and granulocyte (CD66b) lineage surface markers in the neutrophil population in comparison to corresponding isotype controls, determined by flow cytometry. Representative dot plots from three independent experiments (n=3) are shown.

**Table S1**. Set of expressed genes, used for identification of major cell lines in each generated population.

For each population, the four most dominant transcripts (marked in bold), in combination with a q-value < 0.05, were used for assessment of major cell populations. Furthermore, several other dominant transcripts, with a q-value < 0.05 for the corresponding population and a q-value > 0.05 for the other populations are included.

**Table S2**. Significant metabolite changes, with an absolute log_2_ fold change > 1, during erythroid differentiation.

Given are the mean log_2_ fold changes (n=3), compared to the progenitor population and the corresponding FDR adjusted p-value.

**Table S3**. Related to Figure 2a; Significant metabolite changes with an absolute log_2_ fold change > 1, during dendritic cell differentiation.

Given are the mean log_2_ fold changes (n=3), compared to the progenitor population and the corresponding FDR adjusted p-value.

**Table S4**. Significant metabolite changes with an absolute log_2_ fold change > 1, during neutrophil differentiation.

Given are the mean log_2_ fold changes (n=3), compared to the progenitor population and the corresponding FDR adjusted p-value.

**Table S5**. Unique metabolite changes during erythroid differentiation.

Given are the mean log_2_ fold changes (n=3), compared to the progenitor population and the corresponding FDR adjusted p-value.

**Table S6**. Unique metabolite changes during dendritic cell differentiation.

Given is the mean log_2_ fold change (n=3), compared to the progenitor population and the corresponding FDR adjusted p-value.

**Table S7**. Unique metabolite changes during neutrophil differentiation.

Given is the mean log_2_ fold change (n=3), compared to the progenitor population and the corresponding FDR adjusted p-value.

**Table S8**. Common metabolite changes during erythroid and dendritic cell differentiation.

Given are the mean log_2_ fold changes in each population, compared to the progenitor population (n=3). All changes in the erythroid and the dendritic cell population own an FDR adjusted p-value < 0.05.

**Table S9**. Common metabolite changes during erythroid and neutrophil differentiation.

Given are the mean log_2_ fold changes in each population, compared to the progenitor population (n=3). All changes in the erythroid and the neutrophil population own an FDR adjusted p-value < 0.05.

**Table S10**. Common metabolite changes during dendritic cell and neutrophil differentiation.

Given are the mean log_2_ fold changes in each population, compared to the progenitor population (n=3). All changes in the dendritic cell and the neutrophil population own an FDR adjusted p-value < 0.05.

**Table S11**. Common metabolite changes during all differentiations.

Given are the mean log_2_ fold changes in each population, compared to the progenitor population (n=3). All changes own an FDR adjusted p-value < 0.05.

**Table S12**. Excel table containing p-value significant expressional changes in lineages, compared to the progenitor population.

