## Supplemental Tables and Figures for "Metabolite patterns in human myeloid hematopoiesis result from lineage-dependent active metabolic pathways"

^1^Institute of Precision Medicine, Furtwangen University, Jakob-Kienzle-Straße 17, 78054 Villingen-Schwenningen, Germany; ^2^Institute of Pharmaceutical Sciences, University of Freiburg, Albertstraße 25, 79104 Freiburg i. Br., Germany; ^3^CIBSS – Centre for Integrative Biological Signalling Studies, University of Freiburg; ^4^Fraunhofer Institute IZI, Leipzig, EXIM Department, Schillingallee 68, D-18057 Rostock, Germany

***** Corresponding author:**	Prof. Dr. Hans-Peter Deigner,, Phone: +49 7720 307-4232, Fax: +49 7720 307 4725, Institute of Precision Medicine, Furtwangen University, Jakob-Kienzle-Straße 17, 78054 Villingen-Schwenningen, Germany

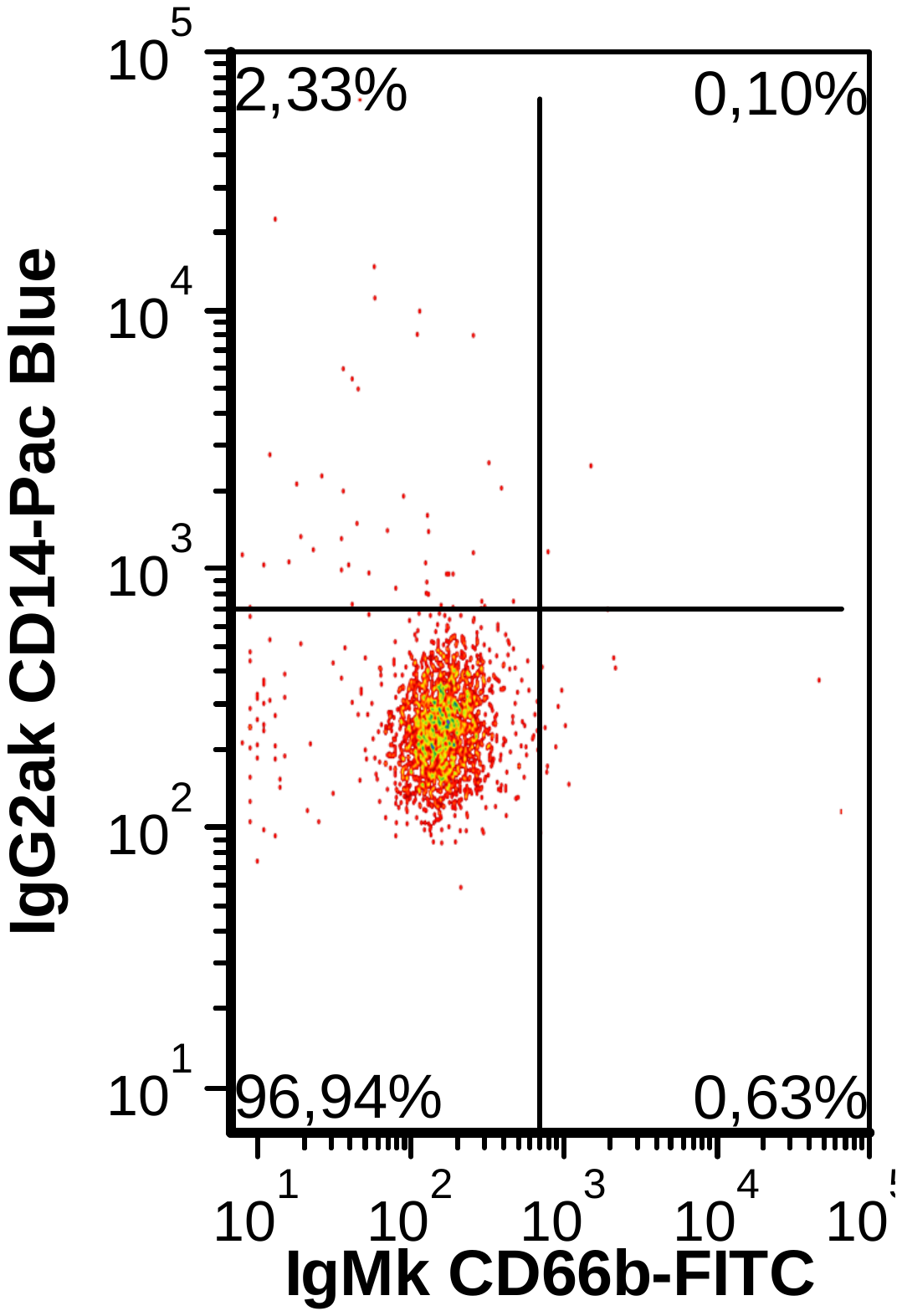

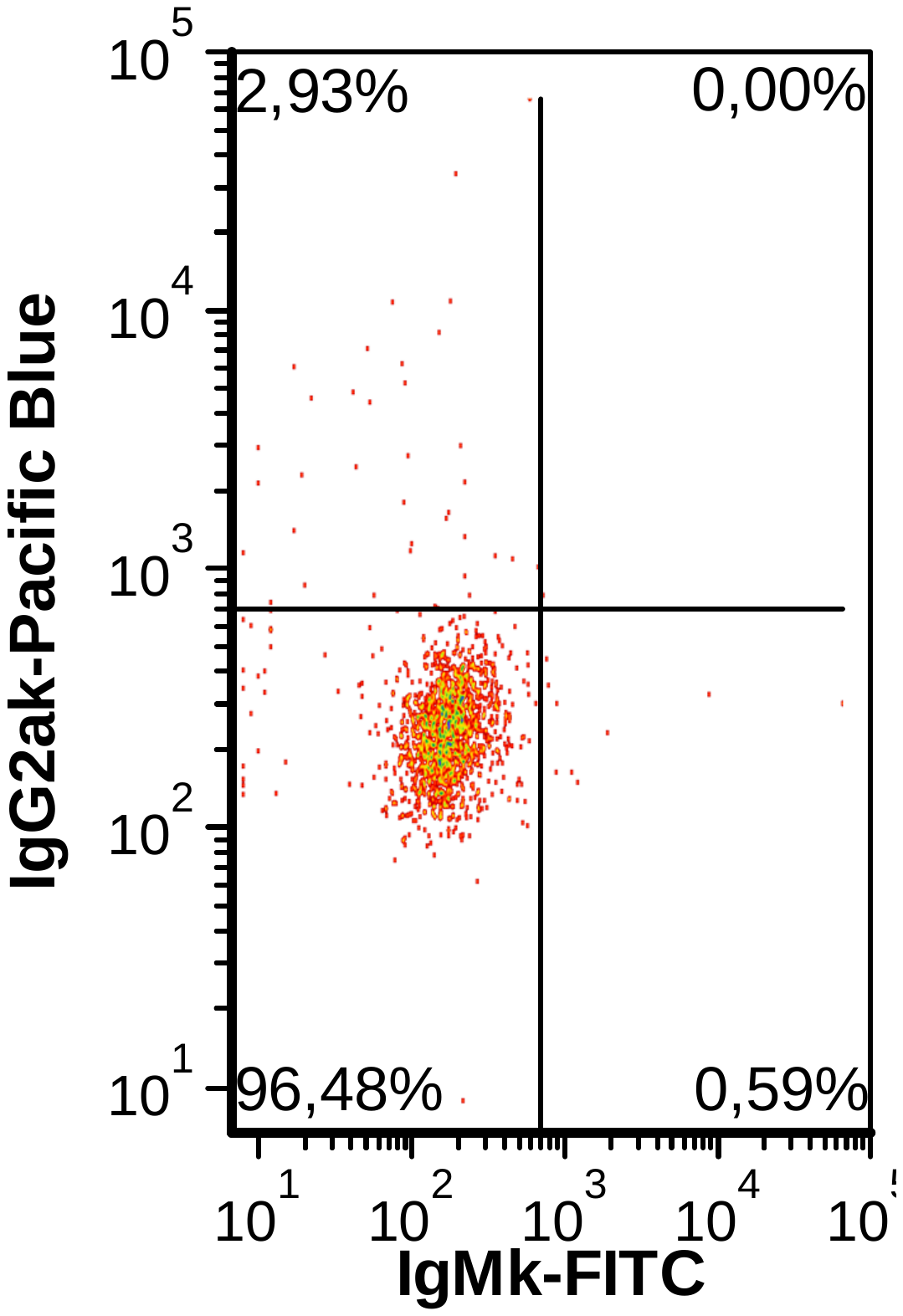

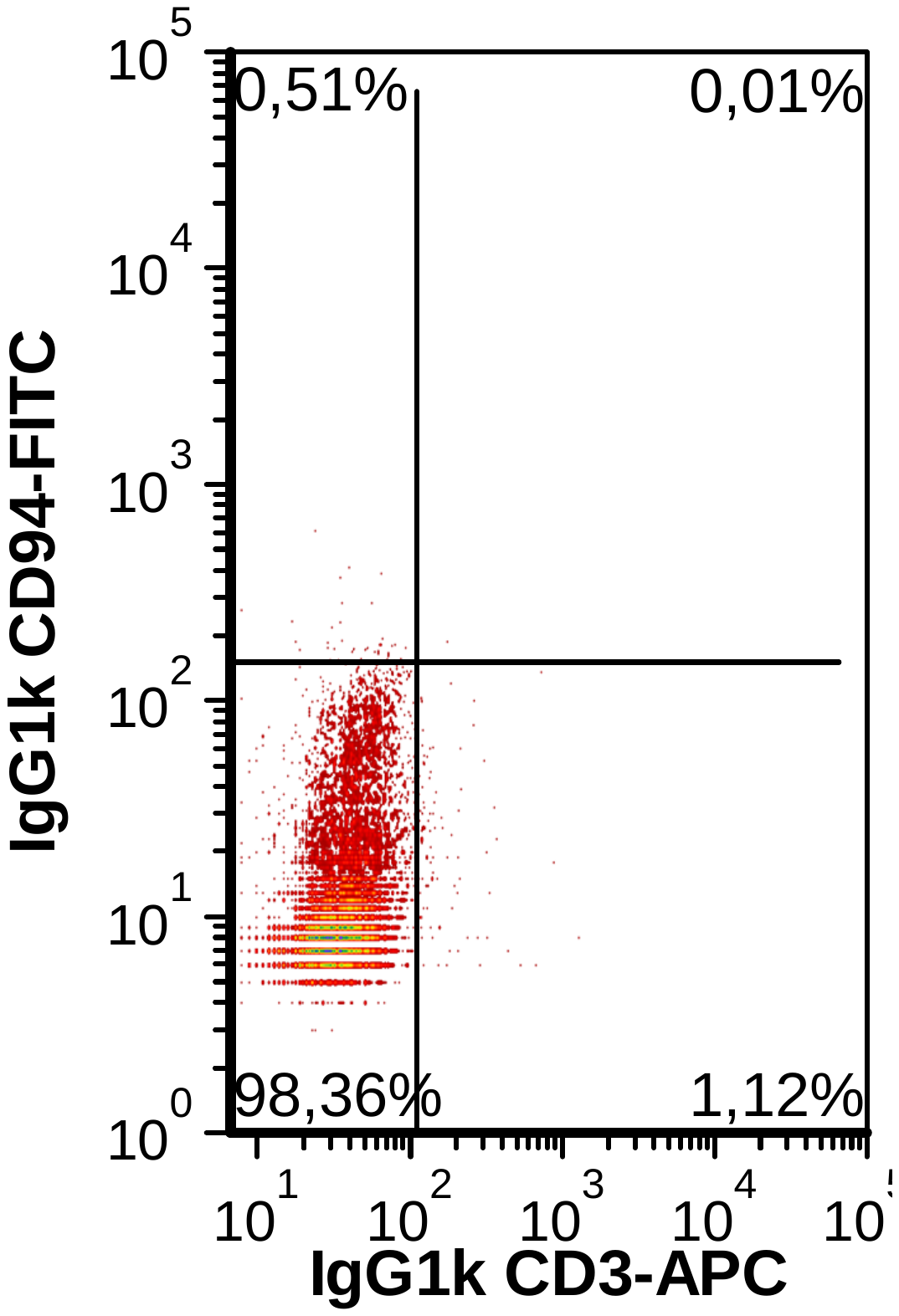

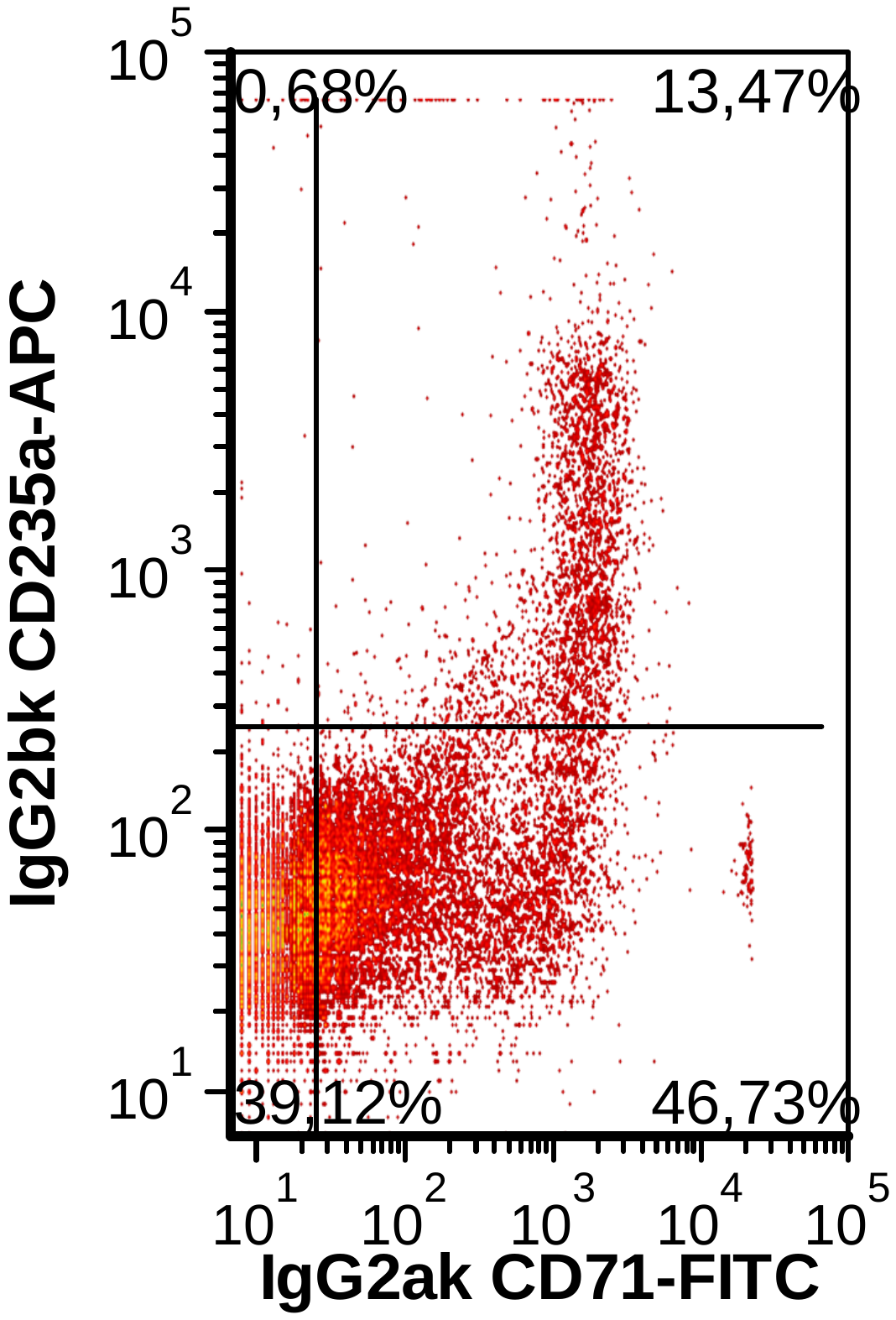

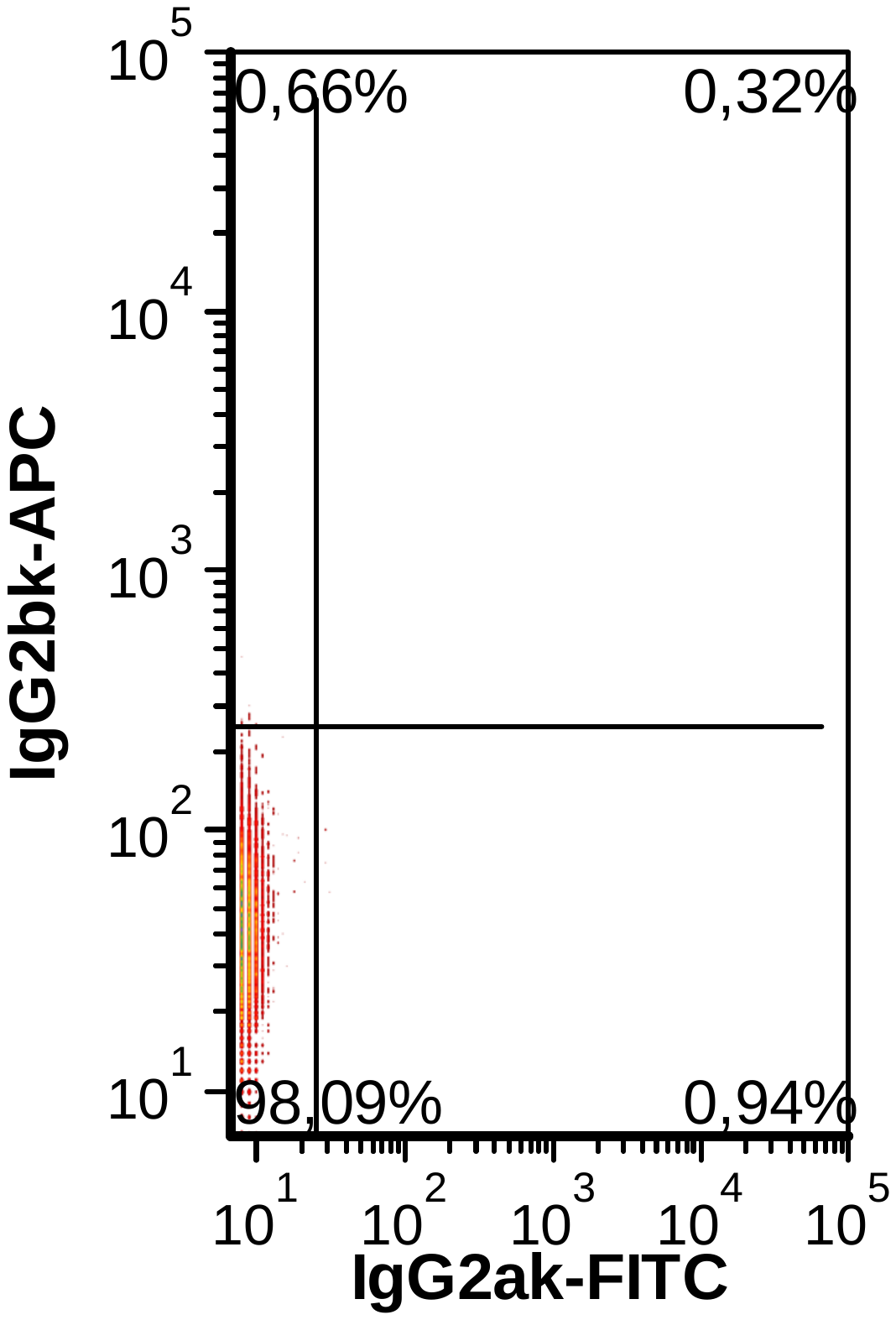

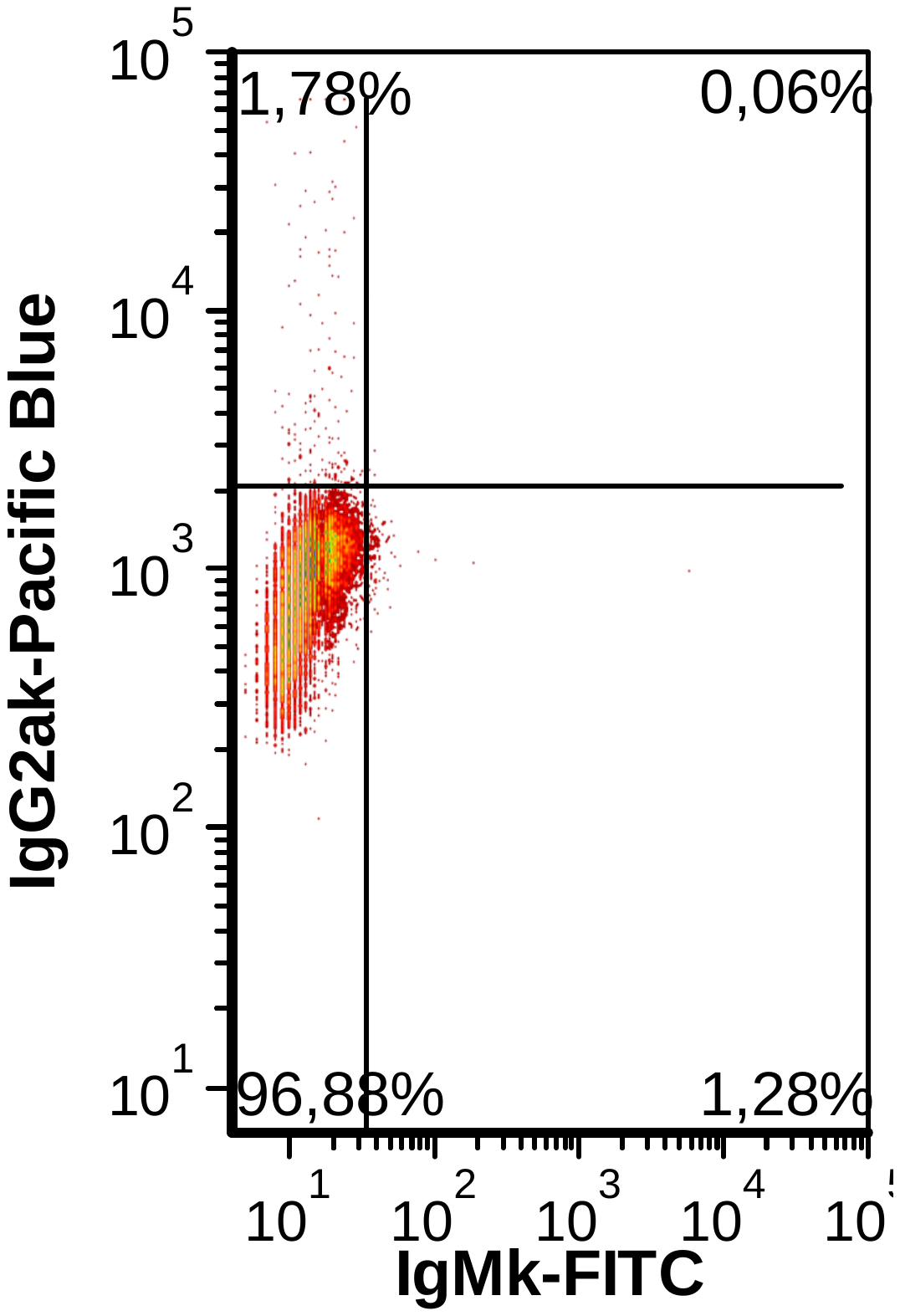

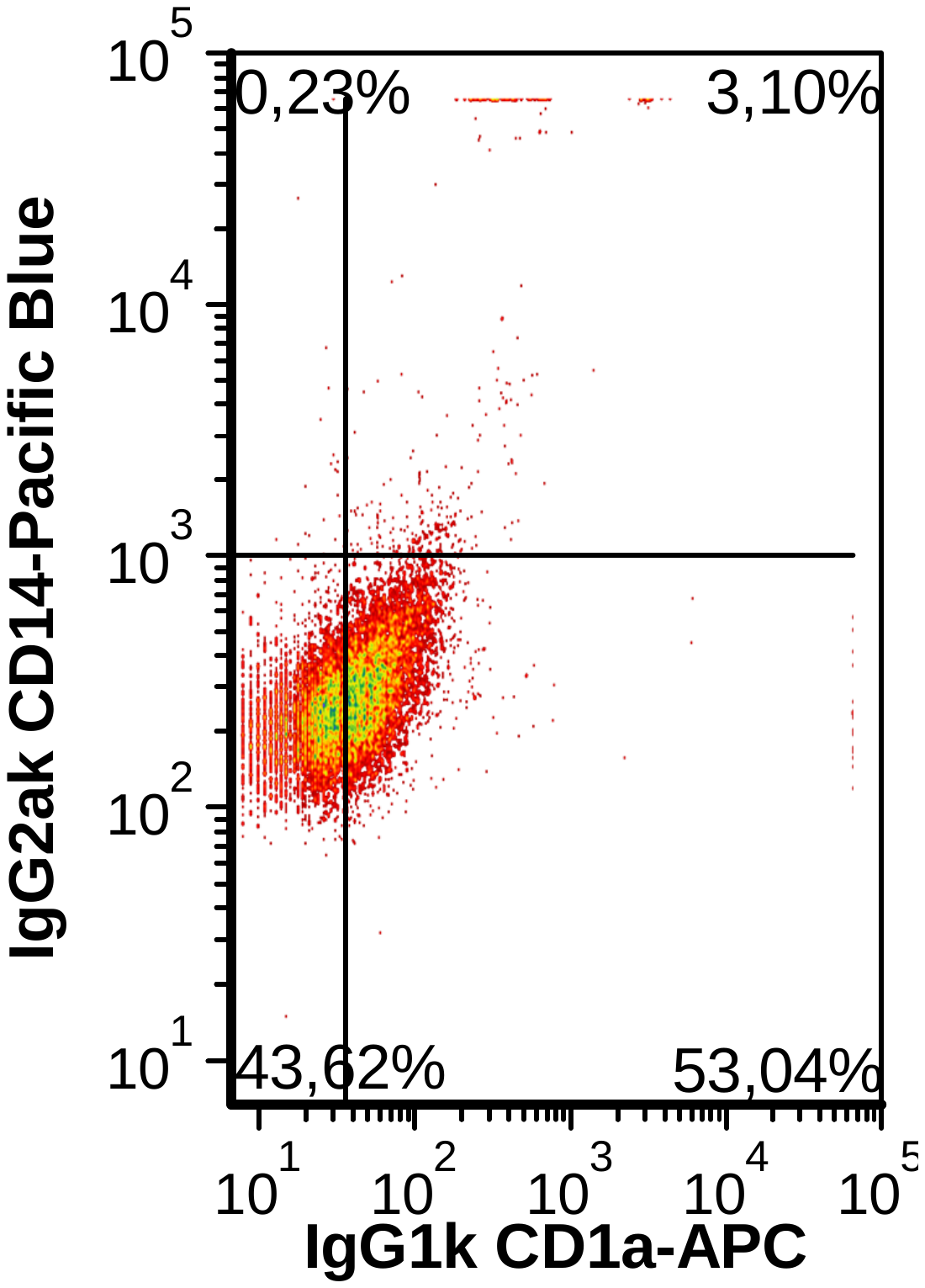

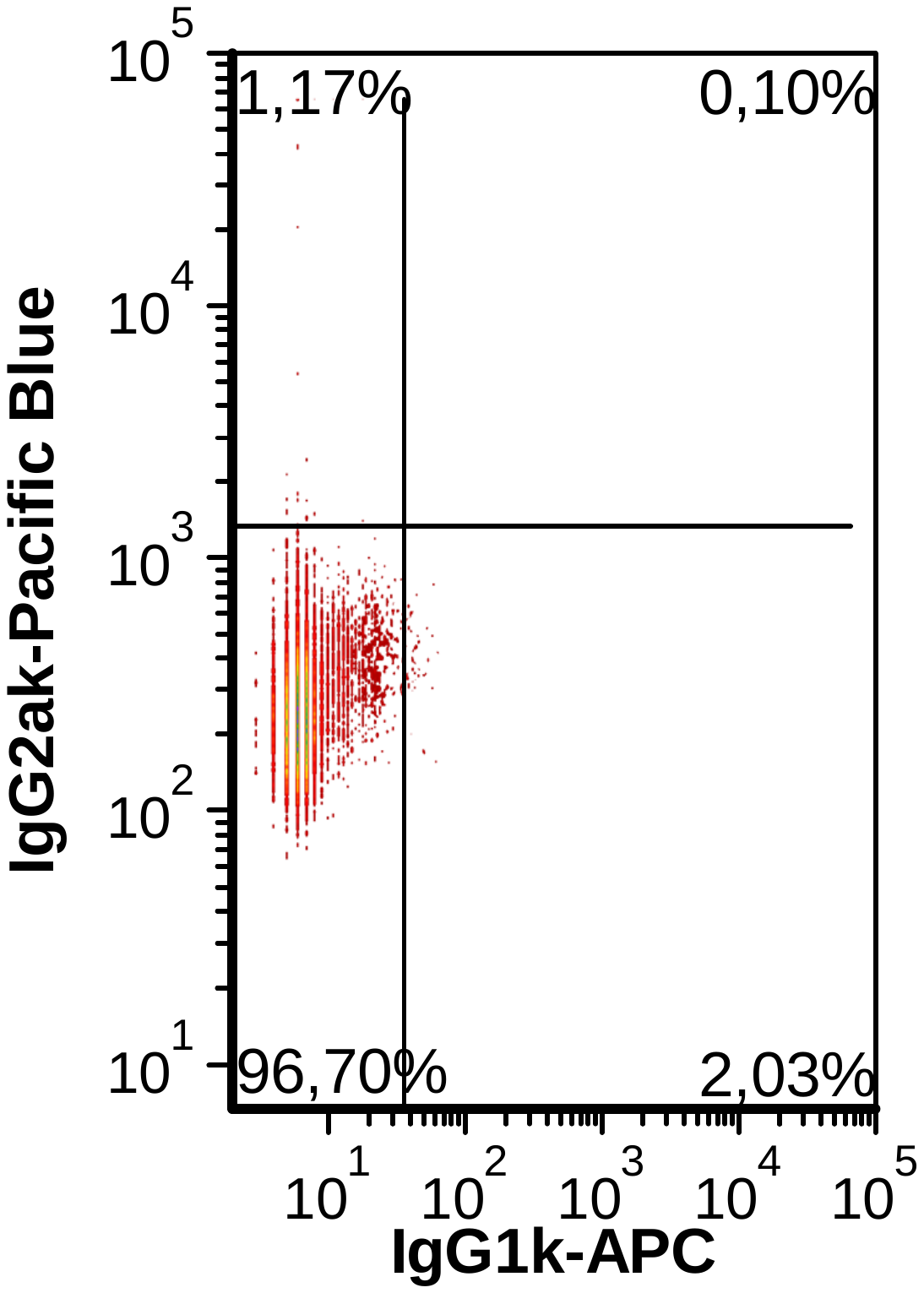

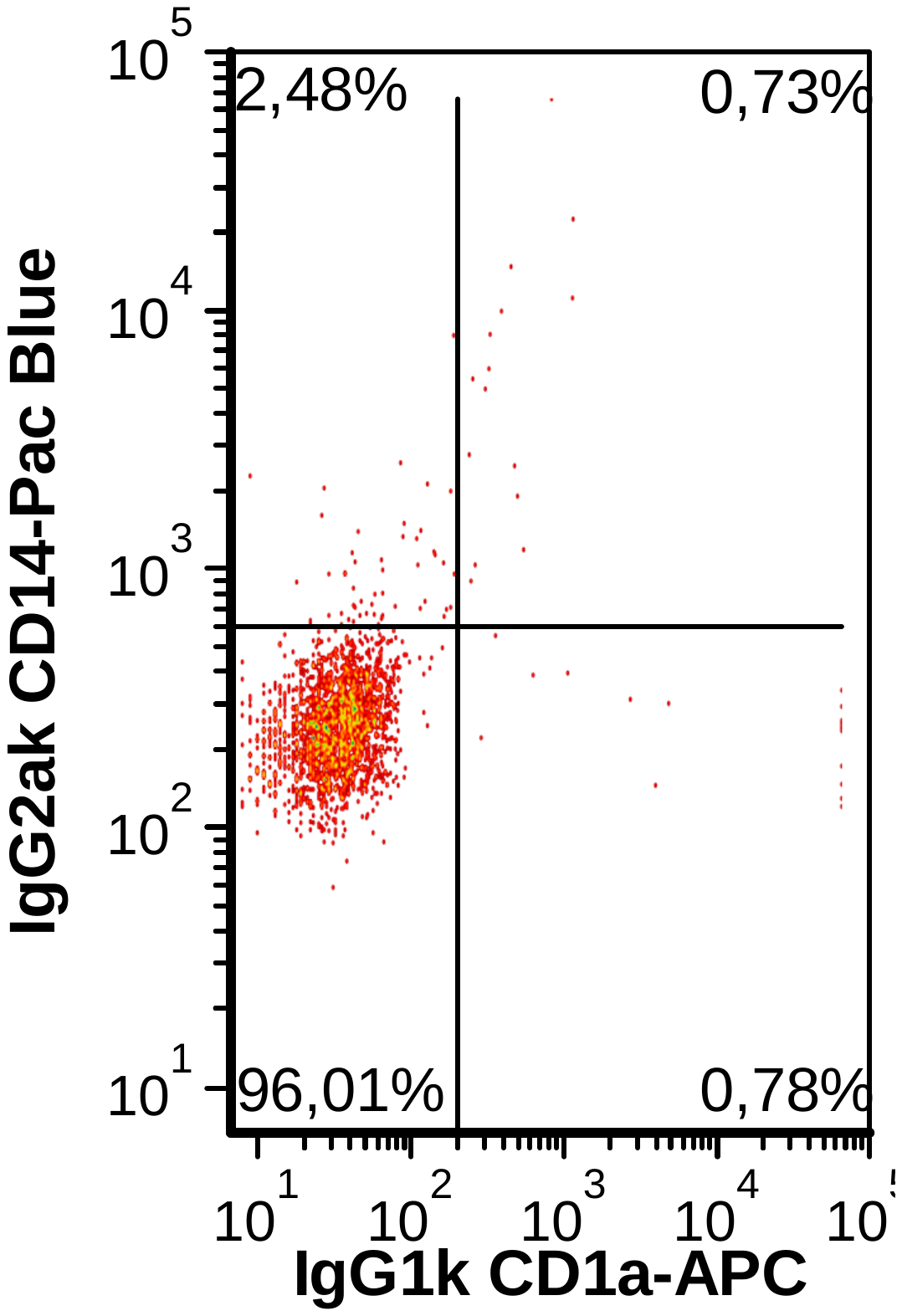

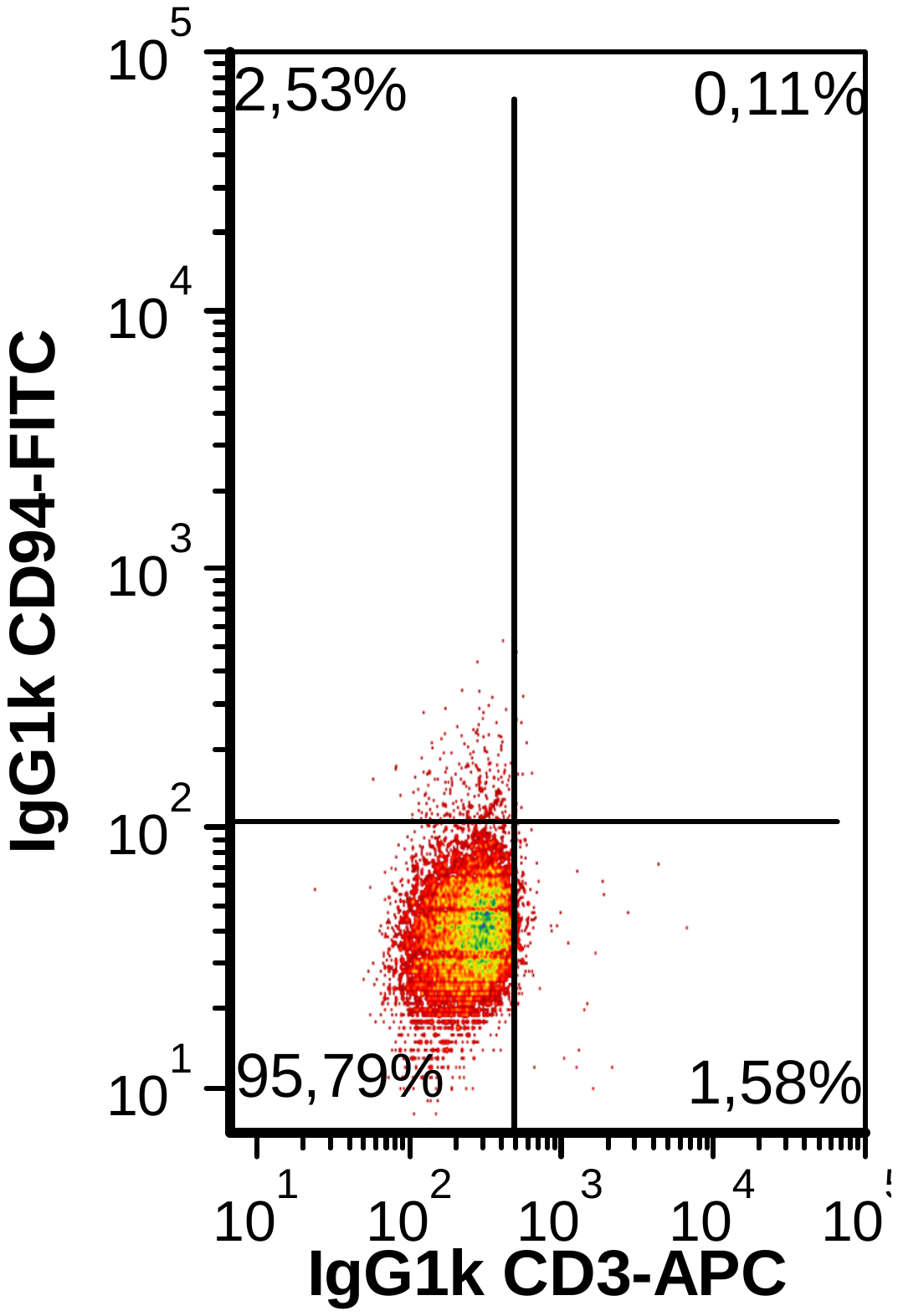

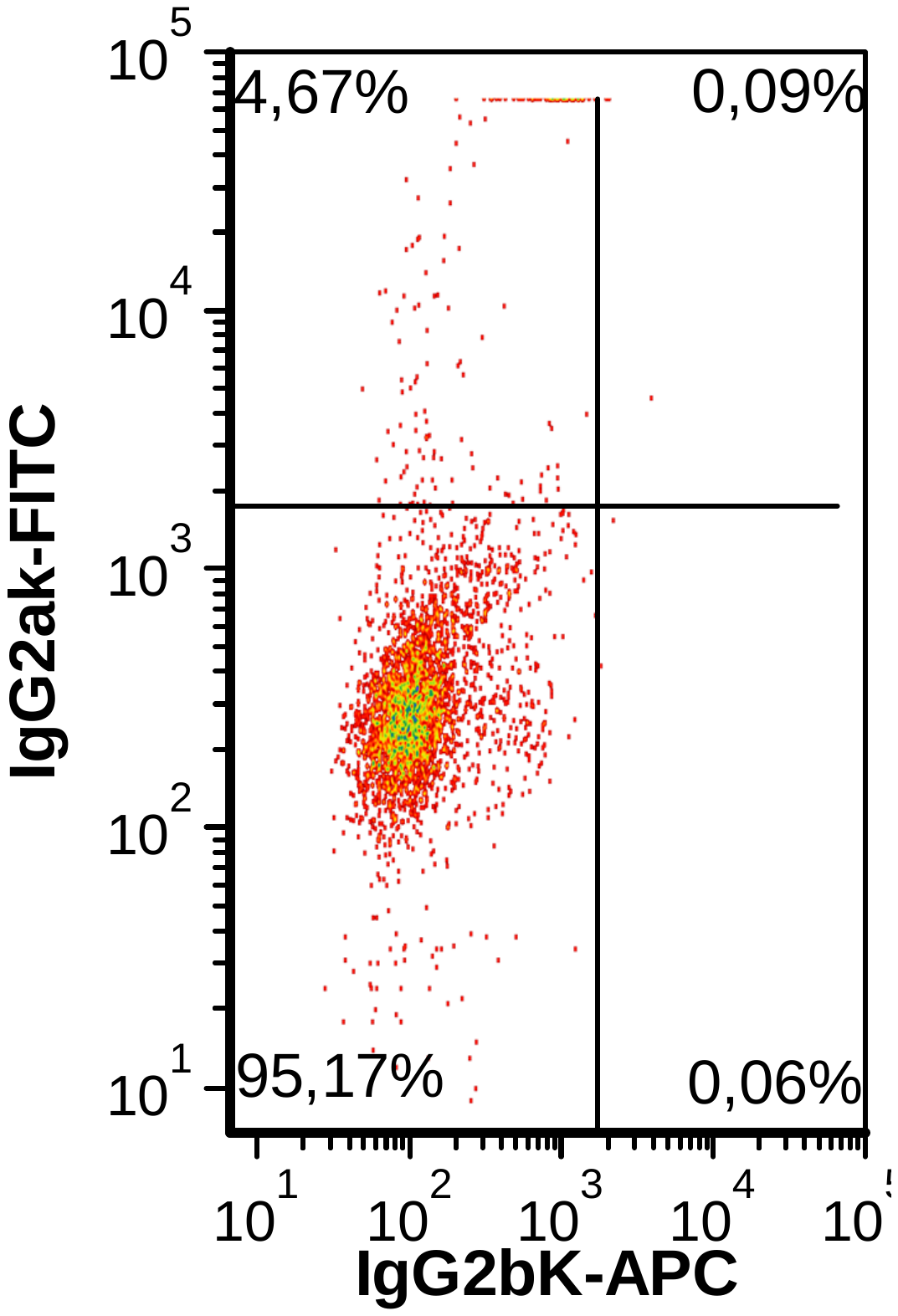

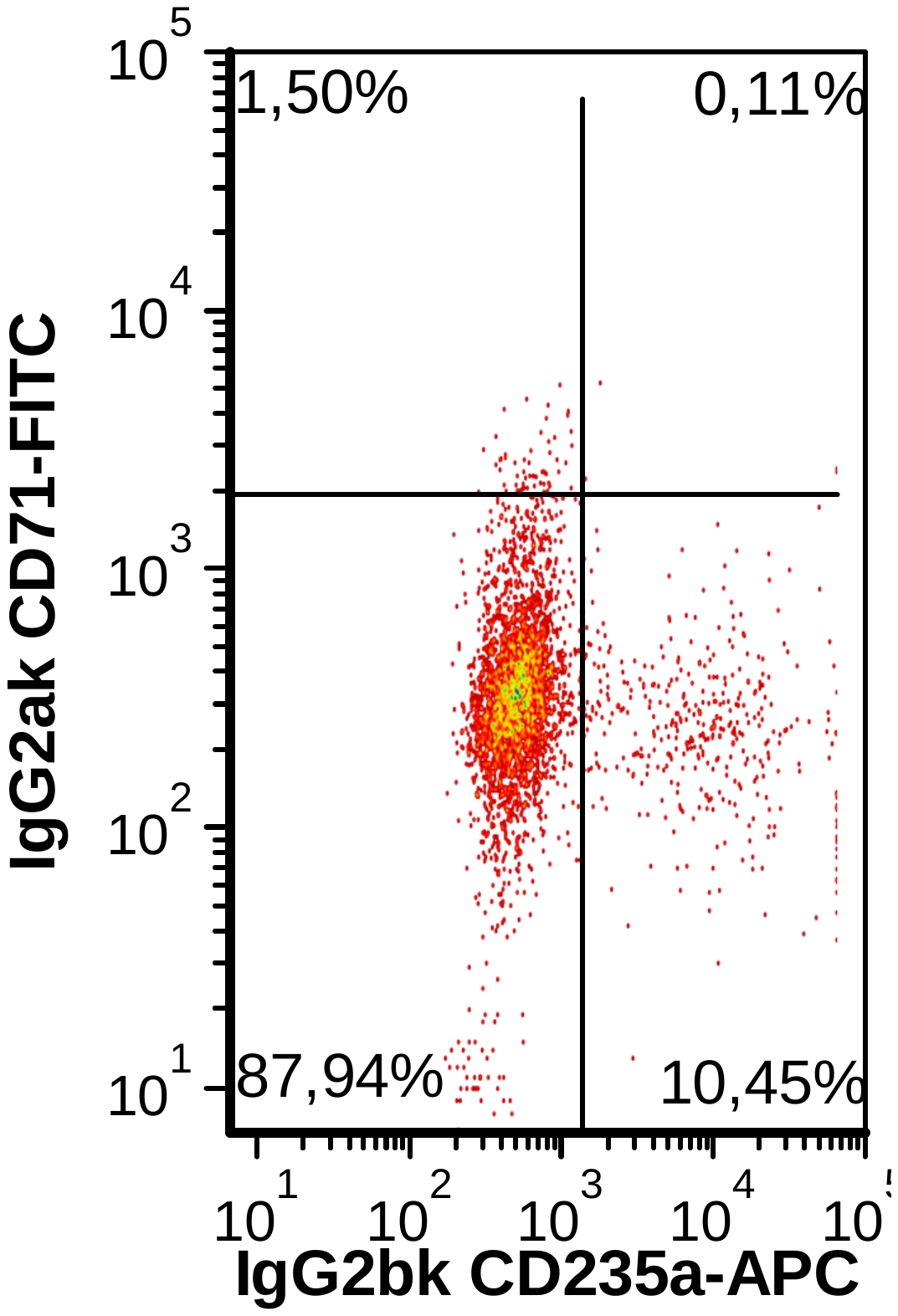

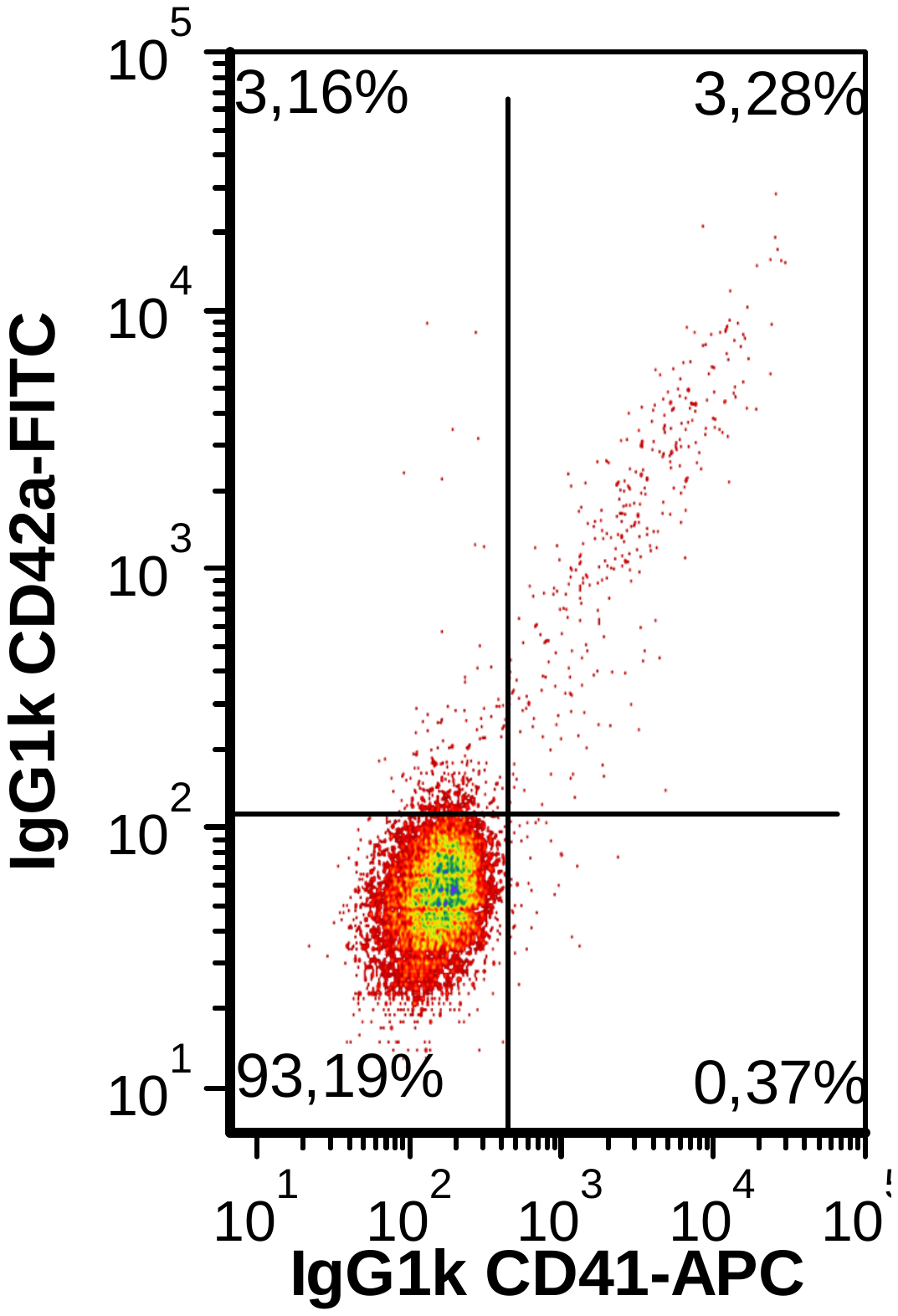

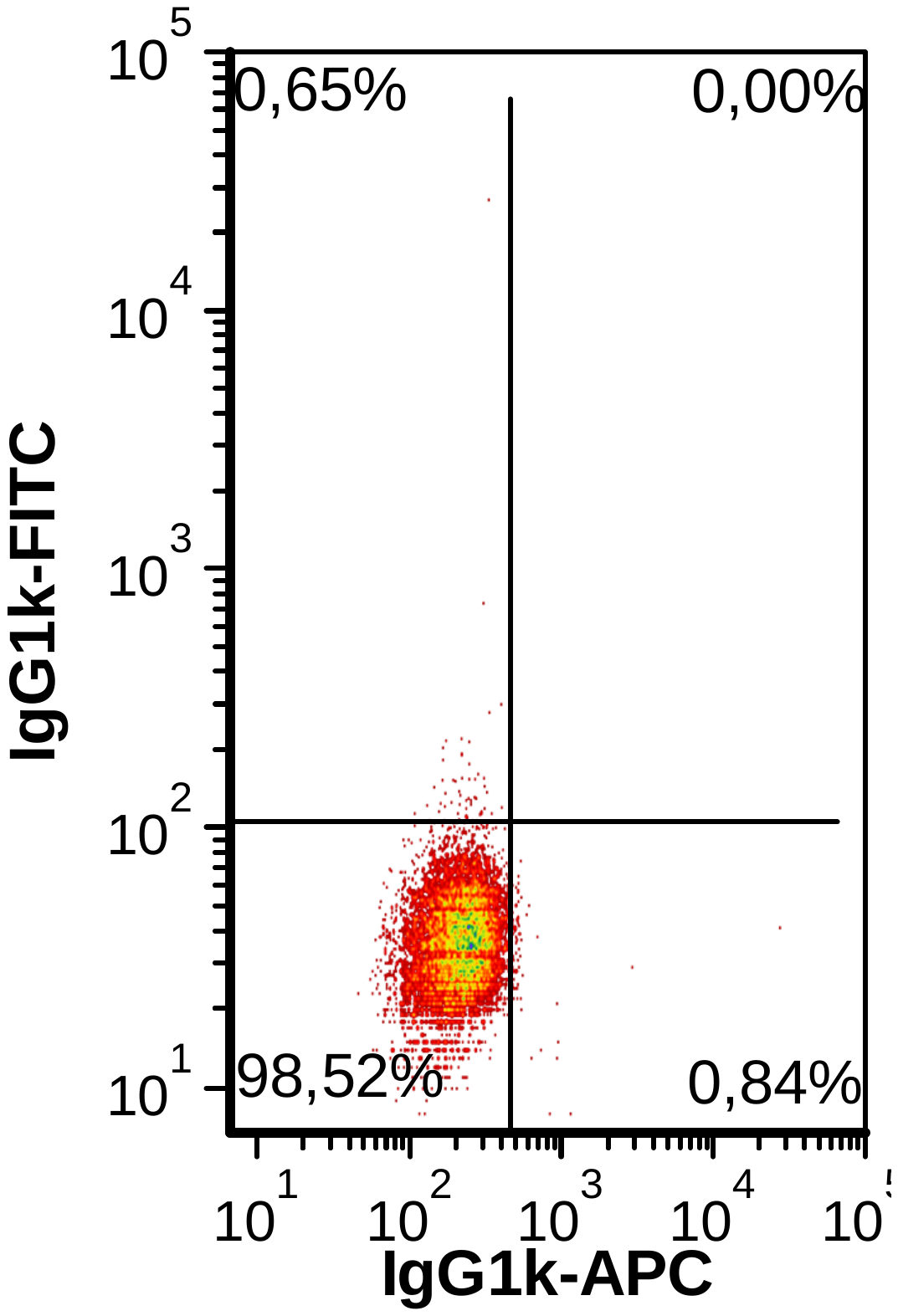

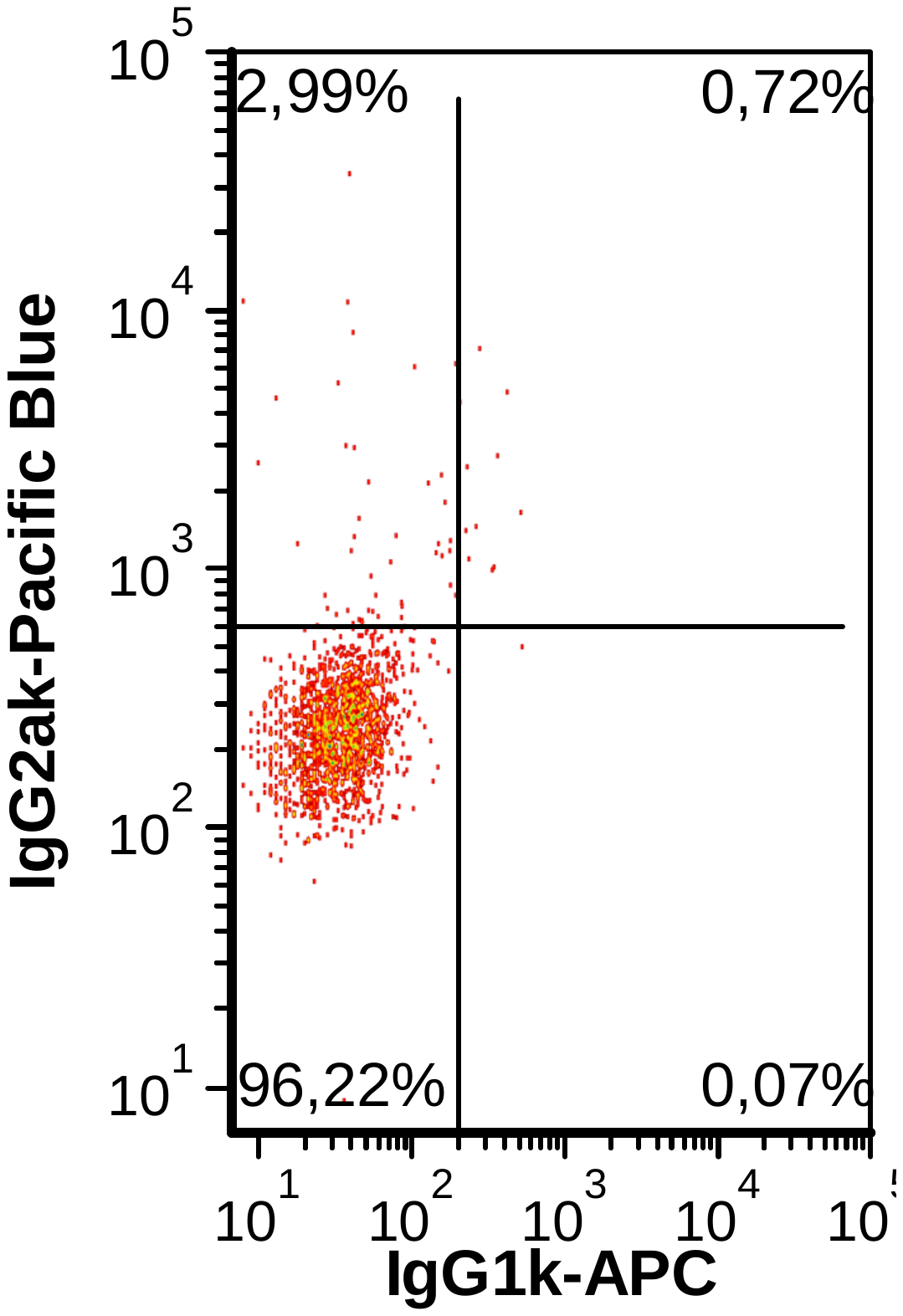

**A**

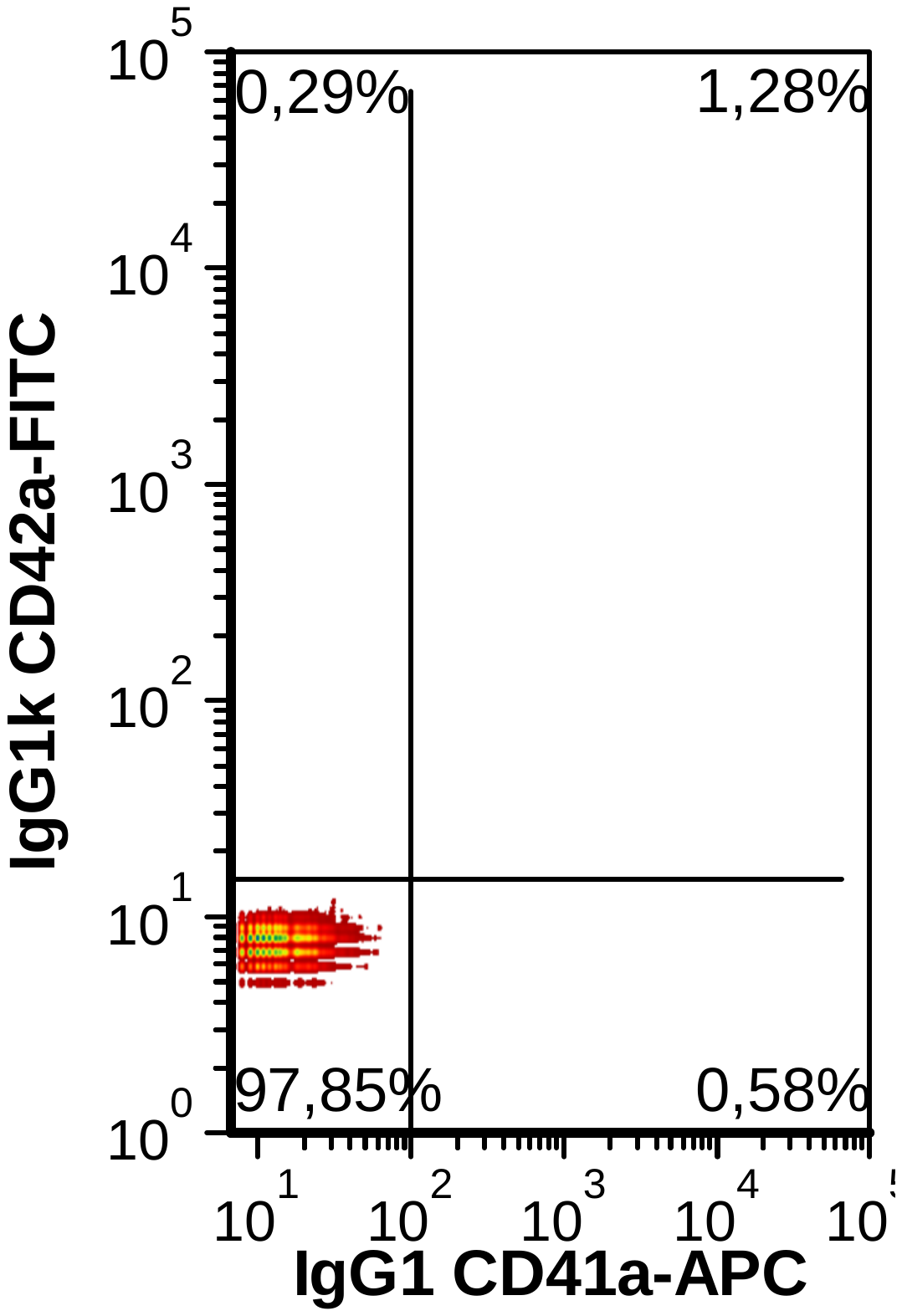

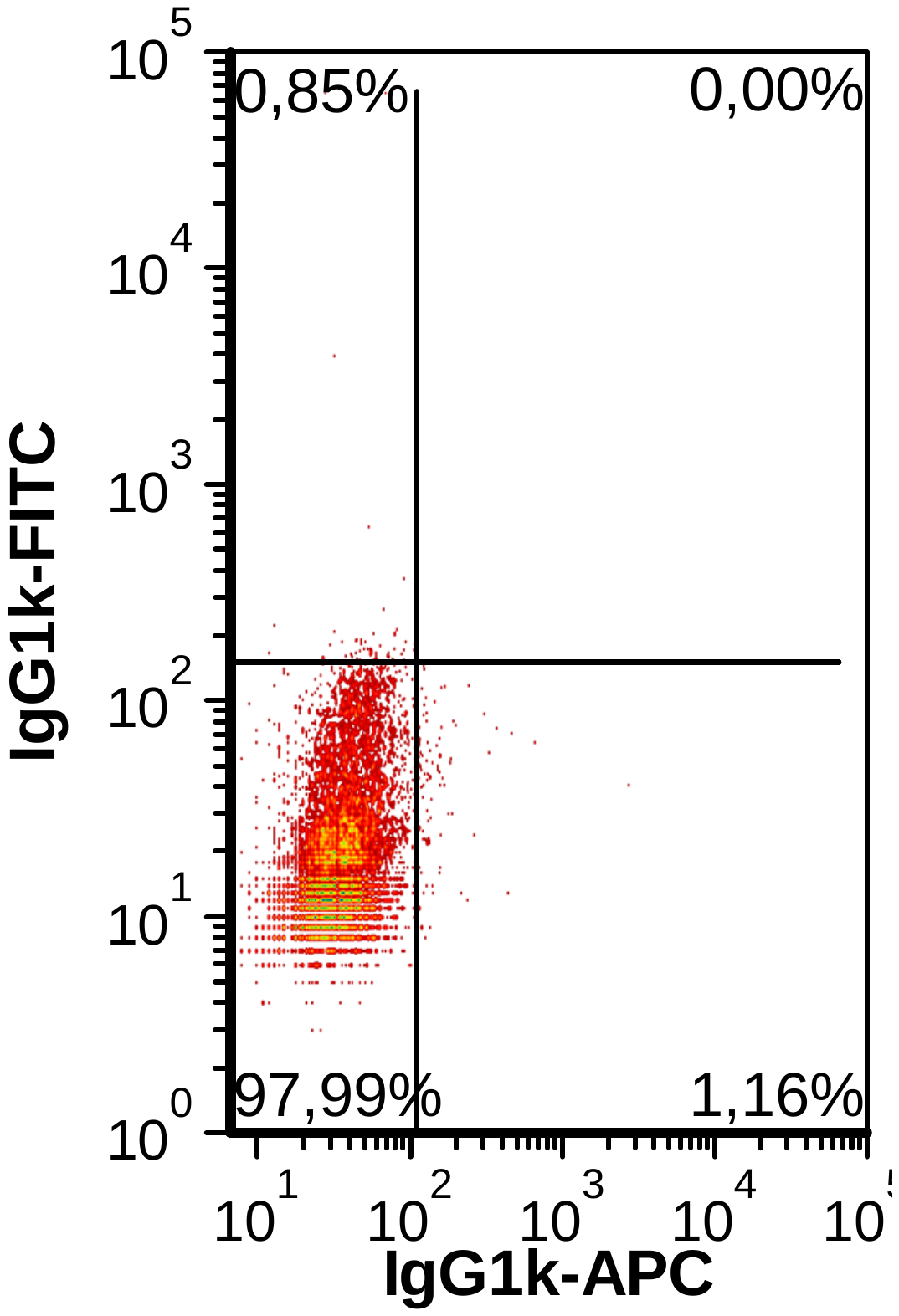

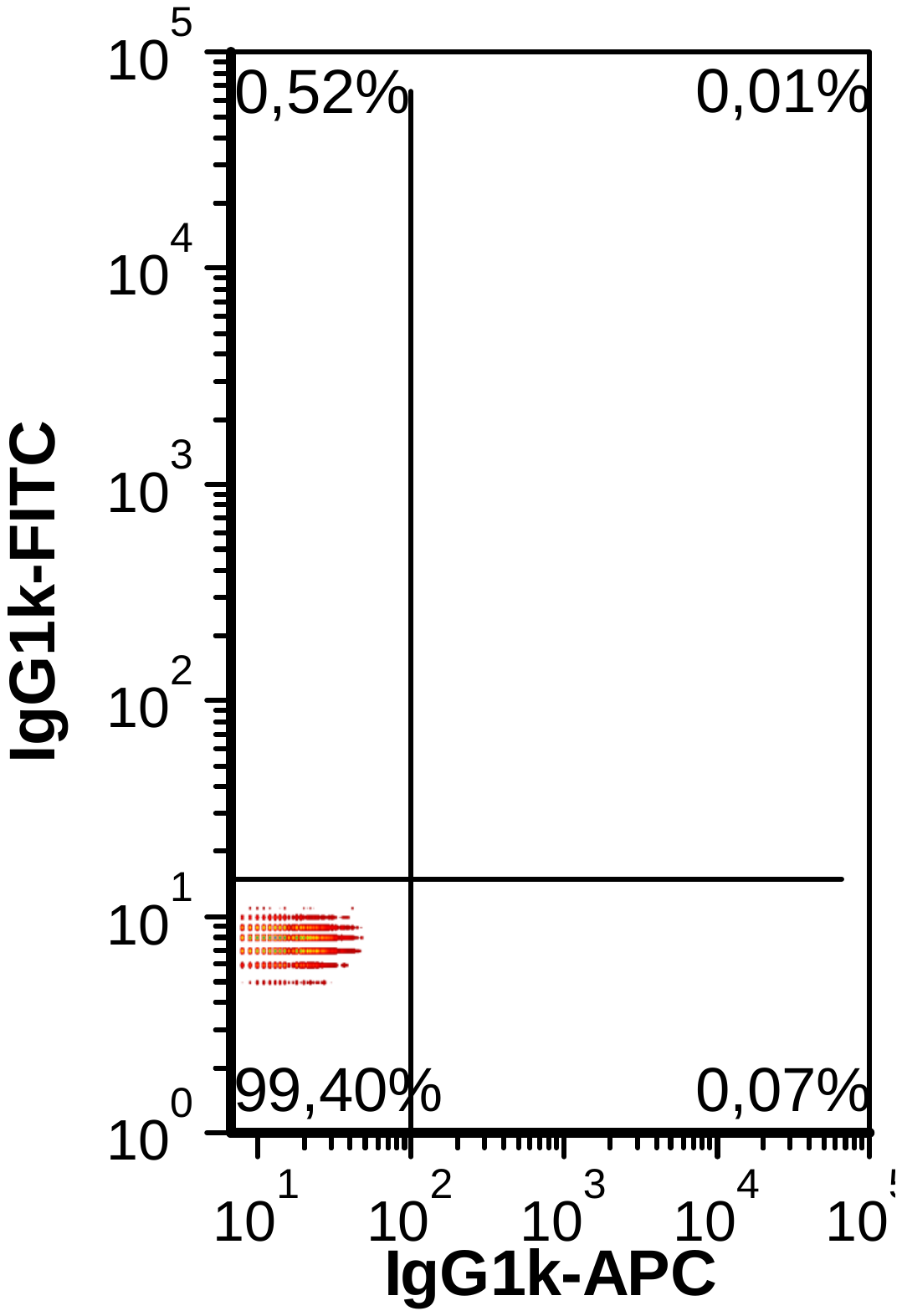

**B**

**C**

**D**

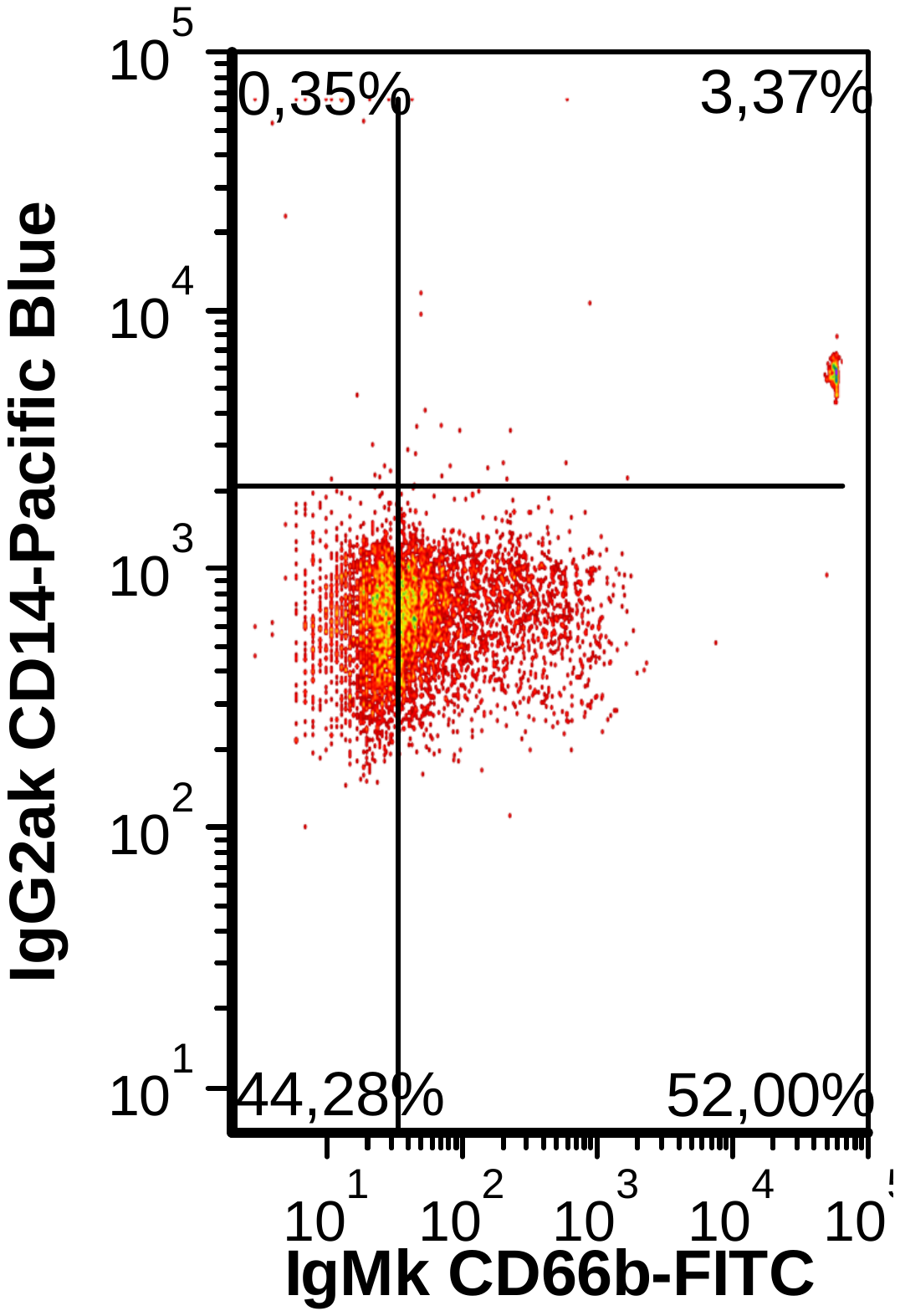

**Figure S1**. Expression of typical lineage markers in the different populations.

(A) Expression of typical lineage selective surface markers in the progenitor population in comparison to corresponding isotype controls, determined by flow cytometry. Representative dot plots from three independent experiments (n=3) are shown.

(B) Expression of erythroid (CD71 and CD235a) lineage surface markers in the erythrocyte population in comparison to corresponding isotype controls, determined by flow cytometry. Representative dot plots from three independent experiments (n=3) are shown.

(C) Expression of megakaryocytic (CD41a, CD42a), monocytic (CD14) and dendritic cell / Langerhans cell (CD1a) lineage surface markers in the dendritic cell population in comparison to corresponding isotype controls, determined by flow cytometry. Representative dot plots from three independent experiments (n=3) are shown.

(D) Expression of natural killer T cell (CD3, CD94), monocytic (CD14) and granulocyte (CD66b) lineage surface markers in the neutrophil population in comparison to corresponding isotype controls, determined by flow cytometry. Representative dot plots from three independent experiments (n=3) are shown.

Table S1. Set of expressed genes in each generated population. For each population, the four most dominant transcripts (marked in bold), in combination with a q-value < 0.05, were used for verification of major cell populations. Furthermore, several other dominant transcripts, with a q-value < 0.05 for the corresponding population and a q-value > 0.05 for the other populations are included.

| **gene** | **Protein** | **Erythrocyte** | **Dendritic cells** | **Neutrophils** | **Q erythrocyte** | **Q dendritic cells** | **Q neutrophils** |
| --- | --- | --- | --- | --- | --- | --- | --- |
| *ALCAM* | CD166 | 23,502 | 108,142 | 21,096 | 0,995 | **0,030** | 0,992 |
| *AZU1* | Azurocidin | 384,931 | 200,726 | **807,309** | 0,026 | 0,127 | 0,026 |
| *BLVRB* | Flavin reductase (NADPH) | 276,353 | 112,461 | 82,272 | **0,026** | 0,989 | 0,992 |
| *C1QA* | Complement C1q subcomponent subunit A | 49,944 | 172,111 | 155,910 | 0,070 | 0,052 | **0,026** |
| *C1QB* | Complement C1q subcomponent subunit B | 44,129 | 249,230 | 226,499 | 0,087 | 0,030 | 0,026 |
| *C1QC* | Complement C1q subcomponent subunit C | 77,604 | **348,741** | 278,409 | 0,026 | 0,030 | 0,043 |
| *CTSG* | Cathepsin G | 126,910 | 73,336 | 313,856 | 0,026 | 0,765 | 0,026 |
| *DPP4* | Dipeptidyl peptidase 4 | 15,285 | 68,023 | 9,535 | 0,087 | **0,030** | 0,581 |
| *ELANE* | Neutrophil elastase | **756,959** | 254,768 | **1721,370** | 0,026 | 0,194 | 0,026 |
| *EPX* | Eosinophil peroxidase | 95,303 | **274,194** | 1,284 | 0,026 | 0,030 | 0,992 |
| *FECH* | Ferrochelatase, mitochondrial | 67,223 | 17,773 | 20,084 | **0,026** | 0,989 | 0,992 |
| *GPNMB* | Glycoprotein NMB | 46.312 | **751.746** | 784.673 | 0.042 | 0.029 | 0.0256 |
| *GP5* | CD42d | 0,000 | 0,000 | 0,000 | 0,026 | 0,030 | 0,058 |
| *HBB* | Hemoglobin subunit beta | **1538,550** | 28,784 | 9,215 | 0,026 | 0,077 | 0,043 |
| *HBM* | Hemoglobin subunit mu | 60,432 | 0,658 | 0,421 | **0,026** | 0,359 | 0,708 |
| *ITGA2B* | CD41 | 38,119 | 21,551 | 34,052 | 0,026 | 0,030 | 0,292 |
| *KEL* | CD238 | 60,424 | 1,476 | 2,070 | **0,026** | 0,503 | 0,992 |
| *KLF1* | Krueppel-like factor 1 | 89,940 | 1,970 | 2,159 | **0,026** | 0,184 | 0,276 |
| *MRC1* | CD206 | 195,641 | **506,575** | 92,780 | 0,026 | 0,030 | 0,992 |
| *PRTN3* | Myeloblastin | **732,813** | 180,812 | **1141,910** | 0,026 | 0,941 | 0,026 |
| *RHAG* | CD241 | 188,878 | 20,615 | 4,118 | **0,026** | 0,989 | 0,344 |
| *S100A8* | Protein S100-A8 | 547,594 | 685,878 | **1475,440** | 0,078 | 0,094 | 0,026 |
| *SLC4A1* | CD233 | 65,298 | 0,250 | 0,353 | **0,026** | 1,000 | 0,992 |
| *TFRC* | CD71 | **490,420** | 131,349 | 85,104 | 0,026 | 0,989 | 0,992 |

Table S2. Significant metabolite changes, with an absolute log_2_ fold change > 1, during erythroid differentiation. Given are the log_2_ fold changes, compared to the progenitor population and the corresponding FDR adjusted p-value.

| **Metabolite** | **logFC** | **adj. p** |
| --- | --- | --- |
| PC aa C42:4 | 1.907 | 0.000 |
| SM-OH C16:1 | 1.802 | 0.002 |
| PC aa C40:3 | 1.769 | 0.000 |
| Histamine | 1.733 | 0.001 |
| PC aa C28:1 | 1.723 | 0.001 |
| lysoPC a C28:0 | 1.681 | 0.001 |
| PC aa C40:2 | 1.656 | 0.000 |
| PC aa C42:5 | 1.611 | 0.000 |
| SM C16:0 | 1.576 | 0.005 |
| PC aa C40:4 | 1.566 | 0.000 |
| PE aa C40:3 | 1.554 | 0.000 |
| PC aa C30:2 | 1.538 | 0.002 |
| SM C18:0 | 1.522 | 0.005 |
| SM C24:1 | 1.515 | 0.007 |
| Putrescine | 1.495 | 0.000 |
| PC aa C34:1 | 1.490 | 0.005 |
| lysoPC a C28:1 | 1.465 | 0.003 |
| SM -OH C14:1 | 1.442 | 0.008 |
| PC aa C32:0 | 1.441 | 0.008 |
| PE aa C40:4 | 1.440 | 0.000 |
| PC ae C40:3 | 1.431 | 0.003 |
| C14:2 | 1.390 | 0.001 |
| C3:1 | 1.385 | 0.000 |
| SM C18:1 | 1.380 | 0.001 |
| PC ae C36:1 | 1.362 | 0.003 |
| PC ae C40:4 | 1.316 | 0.000 |
| C5-DC / C6-OH | 1.308 | 0.001 |
| PC ae C30:1 | 1.306 | 0.005 |
| PE aa C34:1 | 1.296 | 0.000 |
| PC ae C34:0 | 1.291 | 0.013 |
| lysoPC a C24:0 | 1.284 | 0.019 |
| C5:1-DC | 1.280 | 0.000 |
| PS aa C40:3 | 1.273 | 0.013 |
| SM C16:1 | 1.271 | 0.001 |
| PC aa C38:3 | 1.253 | 0.000 |
| PC ae C38:4 | 1.251 | 0.002 |
| C10:1 | 1.250 | 0.005 |
| PC ae C34:1 | 1.246 | 0.012 |
| PC aa C36:1 | 1.241 | 0.018 |
| PE aa C34:0 | 1.236 | 0.000 |
| C12:1 | 1.236 | 0.001 |
| PC aa C42:2 | 1.231 | 0.000 |
| C6 / C4:1-DC | 1.227 | 0.000 |
| PS aa C40:4 | 1.198 | 0.004 |
| C18:1 | 1.194 | 0.000 |
| PC ae C30:0 | 1.190 | 0.002 |
| PC ae C40:5 | 1.190 | 0.000 |
| SM C26:1 | 1.188 | 0.041 |
| PC ae C38:3 | 1.184 | 0.000 |
| PC ae C32:2 | 1.176 | 0.002 |
| C3-DC / C4-OH | 1.175 | 0.003 |
| C18:2 | 1.173 | 0.000 |
| C10 | 1.169 | 0.000 |
| PEA | 1.162 | 0.038 |
| C8 | 1.153 | 0.000 |
| C10:2 | 1.150 | 0.001 |
| C18:1-OH | 1.147 | 0.000 |
| PC aa C42:6 | 1.136 | 0.000 |
| PC aa C32:1 | 1.122 | 0.008 |
| PC ae C36:0 | 1.113 | 0.000 |
| lysoPE a C18:0 | 1.095 | 0.014 |
| PS aa C34:1 | 1.093 | 0.000 |
| PC ae C44:4 | 1.088 | 0.000 |
| C12-DC | 1.082 | 0.000 |
| C9 | 1.081 | 0.002 |
| PC aa C36:6 | 1.081 | 0.000 |
| C5-1 | 1.078 | 0.002 |
| C6-1 | 1.071 | 0.000 |
| PC aa C32:3 | 1.066 | 0.000 |
| PC aa C40:1 | 1.065 | 0.000 |
| PC ae C40:6 | 1.062 | 0.000 |
| C16:1 | 1.058 | 0.000 |
| C7-DC | 1.045 | 0.001 |
| PC ae C32:1 | 1.034 | 0.012 |
| Orn | 1.033 | 0.000 |
| C14 1 | 1.015 | 0.002 |
| C5-OH / C3-DC-M | 1.002 | 0.000 |
| Arg | -1.159 | 0.005 |
| C16:0 Dihydroceramide | -1.793 | 0.001 |
| Ser | -2.094 | 0.001 |
| H1 | -2.483 | 0.000 |

Table S3. Significant metabolite changes with an absolute log_2_ fold change > 1, during dendritic cell differentiation. Given are the log_2_ fold changes, compared to the progenitor population and the corresponding FDR adjusted p-value.

| **Metabolite** | **logFC** | **adj. p** |
| --- | --- | --- |
| Histamine | 2.527 | 0.000 |
| PC ae C30:1 | 2.309 | 0.001 |
| C3:1 | 2.268 | 0.001 |
| SM-OH C14:1 | 2.189 | 0.004 |
| C5-DC C6-OH | 2.177 | 0.004 |
| lysoPC a C28:0 | 2.149 | 0.001 |
| C6 / C4-1-DC | 2.140 | 0.013 |
| SM-OH C16:1 | 2.116 | 0.004 |
| PC ae C32:2 | 2.026 | 0.001 |
| C8 | 1.996 | 0.007 |
| C18:1 Ceramide | 1.975 | 0.000 |
| PC ae C34:1 | 1.970 | 0.008 |
| PC ae C32:1 | 1.961 | 0.004 |
| Taurine | 1.934 | 0.001 |
| C14:1 | 1.915 | 0.003 |
| PC ae C30:2 | 1.899 | 0.000 |
| C6:1 | 1.889 | 0.015 |
| C10 | 1.883 | 0.005 |
| PC ae C34:2 | 1.868 | 0.000 |
| C18:2 | 1.858 | 0.005 |
| PC ae C36:3 | 1.812 | 0.000 |
| C3-DC / C4-OH | 1.797 | 0.004 |
| PC ae C34:3 | 1.768 | 0.000 |
| C7-DC | 1.759 | 0.010 |
| C18:0 Ceramide | 1.744 | 0.000 |
| C16:0 Ceramide | 1.728 | 0.000 |
| C9 | 1.724 | 0.002 |
| lysoPC a C28:1 | 1.705 | 0.003 |
| C10:1 | 1.679 | 0.000 |
| C14:2-OH | 1.669 | 0.003 |
| PC ae C40:3 | 1.652 | 0.004 |
| C5-1-DC | 1.639 | 0.012 |
| PC ae C30:0 | 1.625 | 0.005 |
| SM C16:0 | 1.621 | 0.014 |
| PC ae C38:1 | 1.615 | 0.007 |
| C5:1 | 1.610 | 0.003 |
| C12:1 | 1.591 | 0.000 |
| C4:1 | 1.585 | 0.005 |
| C18 | 1.579 | 0.044 |
| C4 | 1.572 | 0.017 |
| PC aa C28:1 | 1.566 | 0.005 |
| PC ae C36:2 | 1.539 | 0.000 |
| PC ae C40:5 | 1.519 | 0.000 |
| PC ae C38:4 | 1.498 | 0.005 |
| C18:1 | 1.467 | 0.001 |
| C10:2 | 1.460 | 0.000 |
| C14 | 1.438 | 0.017 |
| PC ae C40:2 | 1.399 | 0.013 |
| C16:2 | 1.394 | 0.001 |
| C14:2 | 1.376 | 0.004 |
| C16:2-OH | 1.372 | 0.001 |
| C16:1-OH | 1.366 | 0.003 |
| C3-OH | 1.359 | 0.001 |
| PC ae C40:4 | 1.355 | 0.002 |
| C3 | 1.329 | 0.016 |
| C5 | 1.326 | 0.015 |
| PC ae C40:6 | 1.309 | 0.001 |
| lysoPC a C20:4 | 1.308 | 0.005 |
| C18:1-OH | 1.271 | 0.001 |
| PC ae C36:4 | 1.271 | 0.013 |
| C16-OH | 1.258 | 0.005 |
| PC ae C36:1 | 1.236 | 0.026 |
| C12 | 1.225 | 0.003 |
| PC aa C30:1 | 1.211 | 0.000 |
| PC ae C38:6 | 1.196 | 0.002 |
| PC ae C32:0 | 1.187 | 0.003 |
| C22:0 Ceramide | 1.185 | 0.008 |
| lysoPC a C16:1 | 1.179 | 0.004 |
| PC ae C34:0 | 1.177 | 0.048 |
| C2 | 1.151 | 0.016 |
| SM C18:0 | 1.143 | 0.048 |
| PC ae C44:5 | 1.106 | 0.000 |
| LPE a C22:4 | 1.104 | 0.000 |
| PEA | 1.085 | 0.000 |
| PC ae C44:6 | 1.078 | 0.000 |
| C5-OH / C3-DC-M | 1.051 | 0.000 |
| PC aa C38:0 | 1.049 | 0.004 |
| PS aa C36:2 | 1.045 | 0.000 |
| C14:1-OH | 1.026 | 0.020 |
| SM C26:0 | 1.023 | 0.031 |
| C16:1 | 1.013 | 0.000 |
| C12-DC | 1.001 | 0.000 |
| PC aa C42:4 | -1.005 | 0.017 |
| PE aa C36:4 | -1.061 | 0.000 |
| PE aa C36:5 | -1.063 | 0.000 |
| PE aa C34:3 | -1.086 | 0.000 |
| PC aa C36:4 | -1.123 | 0.007 |
| PE aa C38:7 | -1.129 | 0.001 |
| Arg | -1.342 | 0.001 |
| PC aa C38:5 | -1.470 | 0.000 |
| PC aa C36:5 | -1.471 | 0.000 |
| Ser | -1.547 | 0.006 |
| PE ae C40:1 | -1.548 | 0.000 |
| PE aa C40:7 | -1.552 | 0.000 |
| PE aa C40:6 | -1.557 | 0.000 |
| PC aa C38:6 | -1.656 | 0.000 |
| PE aa C38:6 | -1.750 | 0.000 |
| H1 | -1.864 | 0.000 |
| PE aa C38:5 | -2.021 | 0.000 |

Table S4. Significant metabolite changes with an absolute log_2_ fold change > 1, during neutrophil differentiation. Given are the log_2_ fold changes, compared to the progenitor population and the corresponding FDR adjusted p-value.

| **Metabolite** | **logFC** | **adj. p** |
| --- | --- | --- |
| Histamine | 8.690 | 0.000 |
| Taurine | 5.277 | 0.000 |
| SM -OH C14:1 | 2.821 | 0.001 |
| C2 | 2.498 | 0.000 |
| PC ae C42:4 | 2.373 | 0.006 |
| SM-OH C22:1 | 2.283 | 0.004 |
| SM-OH C22:2 | 2.256 | 0.004 |
| PC ae C38:6 | 2.114 | 0.000 |
| PC aa C38:0 | 1.858 | 0.001 |
| PC ae C40:6 | 1.793 | 0.000 |
| LPE a C22:6 | 1.702 | 0.000 |
| SM-OH C16:1 | 1.483 | 0.043 |
| C3 | 1.474 | 0.014 |
| SM-OH C24:1 | 1.408 | 0.039 |
| PC ae C44:6 | 1.307 | 0.008 |
| PC ae C44:5 | 1.218 | 0.017 |
| PC aa C42:0 | 1.199 | 0.000 |
| PC ae C44:4 | 1.065 | 0.007 |
| PC aa C38:6 | 1.009 | 0.014 |
| PE aa C38:4 | -1.000 | 0.000 |
| PS aa C34:1 | -1.011 | 0.000 |
| PC aa C40:4 | -1.015 | 0.048 |
| PC ae C42:1 | -1.127 | 0.001 |
| PE aa C36:2 | -1.129 | 0.000 |
| PC aa C32:1 | -1.161 | 0.048 |
| PE aa C36:1 | -1.170 | 0.000 |
| PE ae C42:1 | -1.199 | 0.008 |
| PE aa C34:0 | -1.263 | 0.000 |
| PE aa C40:3 | -1.278 | 0.000 |
| PE aa C38:5 | -1.518 | 0.000 |
| PC aa C32:3 | -1.539 | 0.001 |
| Gly | -1.545 | 0.014 |
| PC aa C32:2 | -1.553 | 0.004 |
| SM C22:3 | -1.562 | 0.041 |
| PS aa C40:4 | -1.563 | 0.001 |
| PC aa C36:4 | -1.583 | 0.012 |
| PE aa C34:2 | -1.660 | 0.000 |
| PC aa C30:0 | -1.644 | 0.021 |
| lysoPC a C18:2 | -1.662 | 0.002 |
| PE aa C40:5 | -1.712 | 0.000 |
| PC aa C36:3 | -1.713 | 0.001 |
| PE aa C34:3 | -1.719 | 0.000 |
| Phe | -1.719 | 0.000 |
| Val | -1.785 | 0.038 |
| PC aa C34:4 | -1.791 | 0.000 |
| Tyr | -1.796 | 0.001 |
| PE aa C34:1 | -1.815 | 0.000 |
| H1 | -1.839 | 0.000 |
| Ile | -1.896 | 0.002 |
| Glu | -1.902 | 0.013 |
| PC aa C34:2 | -1.904 | 0.001 |
| PE aa C36:4 | -1.949 | 0.000 |
| Leu | -1.964 | 0.004 |
| Met | -1.978 | 0.002 |
| PC aa C36:2 | -1.984 | 0.001 |
| Thr | -2.002 | 0.012 |
| Asp | -2.008 | 0.014 |
| SM C20:2 | -2.111 | 0.001 |
| PE aa C40:4 | -2.129 | 0.000 |
| C24:1 Ceramide | -2.182 | 0.000 |
| C16:0 Ceramide | -2.264 | 0.000 |
| PC aa C34:3 | -2.269 | 0.000 |
| PE aa C36:3 | -2.461 | 0.000 |
| Pro | -2.692 | 0.000 |
| Lys | -2.823 | 0.000 |
| Ser | -2.841 | 0.000 |
| C18:0 Ceramide | -2.995 | 0.000 |
| His | -3.057 | 0.001 |
| Asn | -3.175 | 0.000 |
| Ala | -3.363 | 0.000 |
| Arg | -3.836 | 0.000 |
| C22:0 Ceramide | -4.024 | 0.000 |
| C16:0 Dihydroceramide | -4.811 | 0.000 |

Table S5. Unique metabolite changes during erythroid differentiation. Given are the log_2_ fold changes, compared to the progenitor population and the corresponding FDR adjusted p-value.

| **Metabolite** | **logFC** | **adj. p** |
| --- | --- | --- |
| Putrescine | 1.495 | 0.000 |
| lysoPC a C24:0 | 1.284 | 0.019 |
| PC aa C32:1 | 1.122 | 0.008 |
| PC aa C32:3 | 1.066 | 0.000 |
| PC aa C38:3 | 1.253 | 0.000 |
| PC aa C40:2 | 1.656 | 0.000 |
| PC aa C40:3 | 1.769 | 0.000 |
| PC aa C40:4 | 1.566 | 0.000 |
| PC aa C42:2 | 1.231 | 0.000 |
| PC aa C42:4 | 1.907 | 0.000 |
| PC aa C42:5 | 1.611 | 0.000 |
| PE aa C34:0 | 1.235 | 0.000 |
| PE aa C34:1 | 1.296 | 0.000 |
| PE aa C40:3 | 1.553 | 0.000 |
| PE aa C40:4 | 1.439 | 0.000 |
| PS aa C40:3 | 1.272 | 0.013 |
| SM C26:1 | 1.188 | 0.041 |

| **Metabolite** | **logFC** | **adj. p** |
| --- | --- | --- |
| C16-OH | 1.258 | 0.005 |
| C16:1-OH | 1.366 | 0.003 |
| PC aa C36:5 | -1.471 | 0.000 |
| PC aa C38:6 | -1.656 | 0.000 |
| PC ae C30:1 | 2.309 | 0.001 |
| PC ae C34:2 | 1.868 | 0.000 |
| PC ae C36:3 | 1.812 | 0.000 |
| PE aa C38:6 | -1.750 | 0.000 |
| PE aa C38:7 | -1.129 | 0.001 |
| PE aa C40:6 | -1.556 | 0.000 |
| PE aa C40:7 | -1.551 | 0.000 |
| C16:0 Ceramide | 1.727 | 0.000 |
| C18:0 Ceramide | 1.743 | 0.000 |
| C18:1 Ceramide | 1.974 | 0.000 |

Table S6. Unique metabolite changes during dendritic cell differentiation. Given is the log_2_ fold change, compared to the progenitor population and the corresponding FDR adjusted p-value.

Table S7Unique metabolite changes during neutrophil differentiation. Given is the log_2_ fold change, compared to the progenitor population and the corresponding FDR adjusted p-value.

| **Metabolite** | **logFC** | **adj. p** |
| --- | --- | --- |
| Ala | -3.363 | 0.000 |
| Arg | -3.836 | 0.000 |
| Asn | -3.175 | 0.000 |
| Asp | -2.008 | 0.014 |
| Glu | -1.902 | 0.013 |
| Gly | -1.545 | 0.014 |
| His | -3.057 | 0.001 |
| Ile | -1.896 | 0.002 |
| Leu | -1.964 | 0.004 |
| Lys | -2.823 | 0.000 |
| Met | -1.978 | 0.002 |
| Phe | -1.719 | 0.000 |
| Pro | -2.692 | 0.000 |
| Thr | -2.002 | 0.012 |
| Tyr | -1.796 | 0.001 |
| Val | -1.785 | 0.038 |
| Histamine | 8.690 | 0.000 |
| Taurine | 5.277 | 0.000 |
| C2 | 2.498 | 0.000 |
| PC aa.C30:0 | -1.644 | 0.021 |
| PC aa.C32:1 | -1.161 | 0.048 |
| PC aa.C32:3 | -1.539 | 0.001 |
| PC aa.C34:2 | -1.904 | 0.001 |
| PC aa.C34:3 | -2.269 | 0.000 |
| PC aa.C34:4 | -1.791 | 0.000 |
| PC aa.C36:2 | -1.984 | 0.001 |
| PC aa.C36:3 | -1.713 | 0.001 |
| PC aa.C38:6 | 1.009 | 0.014 |
| lysoPE a C22:6 | 1.702 | 0.000 |
| PE aa C34:0 | -1.263 | 0.000 |
| PE aa C34:1 | -1.815 | 0.000 |
| PE aa C36:1 | -1.170 | 0.000 |
| PE aa C36:2 | -1.129 | 0.000 |
| PE aa C36:3 | -2.461 | 0.000 |
| PE aa C40:3 | -1.278 | 0.000 |
| PE aa C40:4 | -2.129 | 0.000 |
| PS aa C34:1 | -1.011 | 0.000 |
| PS aa C40:4 | -1.563 | 0.000 |
| C16:0 Ceramide | -2.264 | 0.000 |
| C16:0 Dihydroceramide | -4.811 | 0.000 |
| C18:0 Ceramide | -2.995 | 0.000 |
| C22:0 Ceramide | -4.024 | 0.000 |
| C24:1 Ceramide | -2.182 | 0.000 |
| SM-OH C22:1 | 2.283 | 0.004 |
| SM C20:2 | -2.111 | 0.001 |

Table S8. Common metabolite changes during erythroid and dendritic cell differentiation. Given are the log_2_ fold changes in each population, compared to the progenitor population. All changes in the erythroid and the dendritic cell population own an FDR adjusted p-value < 0.05.

| **Metabolite** | **erythrocytes** | **dendritic cells** | **neutrophils** |
| --- | --- | --- | --- |
| C10 | 1.883 | 1.169 | -0.348 |
| C10:2 | 1.460 | 1.150 | -0.181 |
| C12:1 | 1.591 | 1.236 | 0.075 |
| C12-DC | 1.001 | 1.082 | -0.087 |
| C14:2 | 1.376 | 1.390 | 0.246 |
| C16:1 | 1.013 | 1.058 | 0.010 |
| C18:1 | 1.467 | 1.194 | -0.143 |
| C18:1-OH | 1.271 | 1.147 | -0.060 |
| C18:2 | 1.858 | 1.173 | -0.054 |
| C3:1 | 2.268 | 1.385 | 0.014 |
| C3-DC (C4-OH) | 1.797 | 1.175 | 0.172 |
| C5:1 | 1.610 | 1.078 | -0.100 |
| C5:1-DC | 1.639 | 1.280 | -0.041 |
| C5-DC (C6-OH) | 2.177 | 1.308 | 0.093 |
| C6 (C4:1-DC) | 2.140 | 1.227 | -0.341 |
| C6:1 | 1.889 | 1.071 | -0.121 |
| C8 | 1.996 | 1.153 | -0.316 |
| C9 | 1.724 | 1.081 | 0.033 |
| lysoPC a C28:0 | 2.149 | 1.681 | 0.605 |
| PC aa C28:1 | 1.566 | 1.723 | -0.874 |
| PC ae C30:0 | 1.625 | 1.190 | -0.127 |
| PC ae C34:1 | 1.970 | 1.246 | -0.500 |
| PC ae C38:4 | 1.498 | 1.251 | -0.069 |
| SM C16:0 | 1.621 | 1.576 | -0.344 |
| SM C18:0 | 1.143 | 1.522 | -0.519 |

Table S9. Common metabolite changes during erythroid and neutrophil differentiation. Given are the log_2_ fold changes in each population, compared to the progenitor population. All changes in the erythroid and the neutrophil population own an FDR adjusted p-value < 0.05.

| **Metabolite** | **erythrocytes** | **dendritic cells** | **neutrophils** |
| --- | --- | --- | --- |
| SM-OH C22:2 | 1.577 | 0.298 | 2.256 |
| C16 Dihydroceramide | -1.793 | -0.180 | -4.810 |

Table S10. Common metabolite changes during dendritic cell and neutrophil differentiation. Given are the log_2_ fold changes in each population, compared to the progenitor population. All changes in the dendritic cell and the neutrophil population own an FDR adjusted p-value < 0.05.

| **Metabolite** | **erythrocytes** | **dendritic cells** | **neutrophils** |
| --- | --- | --- | --- |
| Taurine | -0.644 | 1.934 | 5.277 |
| PC aa C36:4 | 0.617 | -1.123 | -1.583 |
| PE aa C34:3 | 0.374 | -1.086 | -1.719 |
| PE aa C36:4 | 0.995 | -1.060 | -1.949 |
| PE aa C38:5 | -0.407 | -2.020 | -1.517 |
| SM C22:3 | 0.763 | -1.240 | -1.562 |

Table S11. Related to Figure 2a; Common metabolite changes during all differentiations. Given are the log_2_ fold changes in each population, compared to the progenitor population. All changes own an FDR adjusted p-value < 0.05.

| **Metabolite** | **erythrocytes** | **dendritic cells** | **neutrophils** |
| --- | --- | --- | --- |
| Arg | -1.158 | -1.341 | -3.836 |
| Ser | -2.093 | -1.547 | -2.841 |
| Histamine | 1.733 | 2.526 | 8.690 |
| PC ae C40:6 | 1.062 | 1.309 | 1.793 |
| PC ae C42:4 | 2.321 | 3.669 | 2.373 |
| SM-OH C14:1 | 1.441 | 2.188 | 2.820 |
| SM-OH C16:1 | 1.801 | 2.1159 | 1.483 |
| H1 | -2.482 | -1.864 | -1.839 |
